# Alternative dimerization is required for activity and inhibition of the HEPN ribonuclease RnlA

**DOI:** 10.1101/2021.03.22.436500

**Authors:** Gabriela Garcia-Rodriguez, Daniel Charlier, Dorien Wilmaerts, Jan Michiels, Remy Loris

**Author notes:** Swiss Federal Institute of Technology Lausanne (EPFL), Lausanne, Switzerland.

## Abstract

The *rnlAB* toxin-antitoxin operon from *Escherichia coli* functions as an anti-phage defense system. RnlA was recently identified as a member of the HEPN (Higher Eukaryotes and Prokaryotes Nucleotide-binding domain) superfamily of ribonucleases. The activity of the toxin RnlA requires tight regulation by the antitoxin RnlB, the mechanism of which remains unknown. Here we show that RnlA exists in an equilibrium between two different homodimer states: an inactive resting state and an active canonical HEPN dimer. Mutants interfering with the transition between states show that canonical HEPN dimerization via the highly conserved RX_4-6_H motif is required for activity. The antitoxin RnlB binds the canonical HEPN dimer conformation, inhibiting RnlA by blocking access to its active site. Single-alanine substitutions mutants of the highly conserved R255, E258, R318 and H323 show that these residues are involved in catalysis and substrate binding and locate the catalytic site near the dimer interface of the canonical HEPN dimer rather than in a groove located between the HEPN domain and the preceding TBP-like domain. Overall, these findings elucidate the structural basis of the activity and inhibition of RnlA and highlight the crucial role of conformational heterogeneity in protein function.

## INTRODUCTION

When a *dmd*^−^ mutant of phage T4 infects *Escherichia coli* cells, gene expression is halted at late stages by the rapid degradation of the phage mRNAs, resulting in the inability of this mutant to proliferate (1). RnlA was first identified as an *E. coli* endoribonuclease responsible for late gene silencing of the *dmd*^−^ mutant T4 phage after infection (2) and later on for post-transcriptionally regulating endogenous *E. coli* gene expression, in particular the synthesis of adenylate cyclase (3). RnlA was subsequently discovered to be the toxin part of the *rnlAB* toxin-antitoxin module, which functions as an anti-phage system in *E. coli* cells, particularly upon infection of the *dmd*^−^ mutant of the T4 phage (4). The endoribonuclease activity of RnlA is controlled by RnlB. RnlA is also directly inhibited by the phage antitoxin encoded by the *dmd* gene (5). The RnlA endoribonuclease constitutes one of the few examples of bacterial toxins with phage-encoded direct inhibitors, which additionally evidences the co-evolution of bacteria and their phages.

RnlA and its plasmid-encoded homolog LsoA have a three-domain structure and a recent bioinformatics analysis suggests that they belong to the HEPN (Higher Eukaryotes and Prokaryotes Nucleotide-binding domain) superfamily, where they define a distinct branch of ribonucleases (6). Their C-terminal domain, previously termed Dmd-binding domain (DBD), contains two conserved arginines and a glutamate (R255, E258 and R318 in RnlA) thought to be active site residues (7–8). The other two domains have identical folds and were termed N-terminal (NTD) and N-repeated (NRD) domain, respectively. The function of the N-terminal domain is unknown, but the N-repeated domain is believed to contribute to substrate binding as the alternative phage-encoded antitoxin Dmd binds in a groove at the interface between the NRD and the HEPN domains close to the location of the above mentioned conserved Arg and Glu residues.

Nucleases are widely represented amongst toxins of Toxin-Antitoxin (TA) systems in bacteria. Endoribonucleases in particular, are most abundant among toxins of type II TA systems (a group of TA system in which both toxin and antitoxin are proteins) (9–10). One of the proposed causes for the extensive prevalence of endoribonucleases as toxins is attributed to the conservation of their RNA targets in all living systems, rendering them effective in a wide range of species in bacteria and archaea, and even in eukaryotes. Endoribonucleases from TA systems have been successfully employed as tools for genetic manipulation of yeasts (11), in the cell-ablation-based bio-containment strategy in *Arabidopsis thaliana* (12), and in the development of antiviral and anticancer gene therapy strategies (13–14).

The HEPN domain was first described as the 110 residues-long domain on the C-terminus of human protein Sacsin. HEPN domains are widely distributed amongst many bacterial and archaeal species, and also amongst animals (15). The HEPN domain consists of five α-helices, with three of them arranged in an up-and-down helical bundle and two shorter helices on one side (15). According to their domain architectures, gene-neighborhoods, phyletic patterns and operon organization, seven distinctive HEPN domain families can be discerned. Many function as metal-independent RNases that take part in cellular RNA maturation as well as in biological conflicts, such as virus-host interactions and as part of selfish genetic elements (6). All these HEPN proteins contain a RX_4-6_H motif as their most strongly conserved feature (where X represents any residue). The Arg and His in this motif were shown to be essential for the endoribonuclease activity in a number of widely divergent HEPN proteins (16–17).

When HEPN domains were first identified in bacteria and archaea, they were found to frequently co-localize with genes encoding minimal nucleotidyltransferases (MNTs) (15), which suggested the concerted action of both gene products. Interestingly, the RX_4-6_H motif is usually lost in HEPN-MNT fused aminoglycoside NTases (18–19) (e.g., PDB entries 1KNY and 4CS6), where the HEPN domain is believed to have preserved a nucleotide binding function upon the loss of its RNase activity (20). However, non-covalent HEPN-MNT pairs have since then been shown to form typical type II TA systems, e.g. in *Shewanella oneidensis* (21–22) where the HEPN toxin functions as an RNase that is directly inhibited by its cognate MNT antitoxin. HEPN-MNT TA modules are since then identified as the most widely distributed class of mobile two-gene modules in prokaryotes, but their functions and mechanisms remain elusive (23).

Here, we analyze the functional mechanism of the *E. coli* toxin RnlA. We show that, in contrast to what was suggested in earlier work (7), the dimer observed in the crystal structure of the free toxin corresponds to a non-active precatalytic conformation. Structural and mutagenesis studies show that RNase activity requires the formation of a canonical HEPN dimer and that the active site is located near this HEPN dimer interface and not at the interface between the N-repeated domain and the HEPN domain previously observed in the crystal structure of the free RnlA dimer (7–8).

## MATERIALS AND METHODS

### Cloning, expression and purification of His-RnlA and RnlA-RnlB-His

An expression clone for wild-type His-RnlA (*his-rnlAB*_pET28a) and the RnlA:RnlB-His complex (*rnlAB-his*_pET21b) in BL21(DE3)) was available from previous work (24). Site-directed mutagenesis of RnlA was carried out by PCR extension or full-plasmid amplification with partially overlapping primers (25) containing a single amino acid mutation, from the His-RnlA expressing plasmid *his-rnlAB*_pET28a. Previously the *rnlA* coding region was amplified by PCR to exchange the initial thrombin cleavage site between the N-terminus His-tag and the start of the *rnlA* gene for the TEV cleavage sequence. The resulting genes encoding the N-terminal His-tagged R255A, E258A, R318A, H323A, V206R and D245R RnlA mutants were cloned into pET28a plasmids under the control of the inducible T7 promoter and transformed into BL21(DE3) competent cells for further protein expression and purification. The RnlA truncation was generated by NEB Assembly from the *his-rnlAB_pET28a* plasmid. The resulting genes encoding the N-terminal His-tagged RnlAΔ1-91 in presence and absence of RnlB were cloned into a *pET21b* vector. Each construct was verified by DNA sequencing.

RnlA wild-type and RnlA:RnlB-His complex were produced and purified as described by Garcia-Rodriguez and co-workers (24). The endogenous *rnlAB* operon is expressed under the control of the inducible T7 promoter in BL21(DE3) cells. Placement of the affinity tag on RnlA leads to loss of RnlB antitoxin during Ni-NTA purification followed by gel filtration. A homogeneous preparation of RnlA:RnlB containing stoichiometric amounts of both proteins could only be obtained using a His tag placed C-terminal to RnlB.

For RnlA mutants and the RnlAΔ1-91 (TBP2-HEPN) truncated version, BL21(DE3) cells containing the pET28a *rnlA_mutant* or the *RnlAΔ1-91-rnlB* pET21b plasmids were grown overnight in LB medium supplemented with 50 μg ml^−1^ kanamycin or 100 μg ml^−1^ ampicillin, respectively, at 310 K. A 100-fold dilution was used to inoculate 1 l cultures of the same medium that were incubated at 310 K until OD_*600*_ reached 0.6, then the temperature was decreased to 293 K and expression of the desired proteins was induced with 0.5 mM isopropyl-β-D-thiogalactopyranoside (IPTG). Cells were harvested after 20 hours of expression by centrifugation at 277 K and the pellets were resuspended in lysis buffer (20 mM Tris-HCl, pH 8, 500 mM NaCl, 2 mM 2-mercaptoethanol) supplemented with a protease inhibitor cocktail (cOmplete, Sigma-Aldrich/Merck). Cell extracts were injected into a Ni Sepharose column for affinity chromatography purification followed by size exclusion chromatography of the RnlA-mutant–containing fractions as a final step. The progress of the purification was verified by SDS-PAGE (26) and anti-Histidine tag Western blot (27).

### Cell-free expression of RnlB antitoxin

Successful expression and purification of RnlB in *E. coli* failed and therefore synthesis of C-terminally His-tagged RnlB (RnlB-His) in a cell-free expression system was carried out using WEPRO®7240H Expression Kit (CellFree Sciences Co., Ltd). The *rnlB* gene with a C-terminal His-tag sequence was cloned into the pEU-E01 vector under the SP6 promoter. The final *rnlB-his-*pEU-E01 plasmid was purified, verified by DNA sequencing and subsequently used for “large” scale transcription and translation in reaction volumes of 250 μl and 501 μl, respectively, according to the kit manufacturer’s instructions. To isolate RnlB-His from the components of the wheat germ extract expression kit, a Ni-NTA purification in batch mode was carried out. 100 μl of pre-washed Ni Sepharose High Performance resin (GE Healthcare) was added to the final translation mixture including WEPRO®7240H. After 15 minutes incubation at room temperature on a shaking platform, the unbound fraction was separated by centrifugation and the Ni-bound proteins were washed two times with 20 mM Tris-HCl, pH 8, 500 mM NaCl, 1 mM tris(2-carboxyethyl) phosphine. RnlB-His bound to the Ni beads was eluted by adding 500 mM imidazole to the wash buffer. The eluted fraction was then injected onto a Superdex column for final purification. The progress was monitored by SDS-PAGE and anti-Histidine Western blot.

### Ribonuclease activity assays

RNase activity of RnlA and its mutants was monitored by following the cleavage of bacteriophage MS2 genomic RNA (Roche Diagnostics) via denaturing gel electrophoresis as described (28). RNase-free purifications of all proteins were completed before carrying out the activity assays, by preparing buffers with diethyl pyrocarbonate (DEPC)-pretreated water and/or Ultrapure RNase-free water, using RNase-free laboratory consumables and RNase-cleaned surfaces. 50 pmol of RnlA, the reconstituted RnlA:RnlB complex at a 1:2 molar ratio, and 50 pmol of the four alanine substitution RnlA mutants, were all incubated with 0.08 μg μl^−1^ of MS2 RNA in 10 μl final volume, in the presence of RNase Inhibitor (SUPERase In™ RNase Inhibitor, Invitrogen). All reactions were incubated for 1 h at 310 K in PBS. The reactions were stopped by addition of 10 μl of 2x loading dye (95 % v/v formamide, 0.02 % m/V SDS, 0.02 % m/V bromophenol blue, 0.01 % m/V xylene cyanol, 1 mM EDTA). Samples were loaded after 10 minutes incubation at 343 K onto 6 % urea gels which had been pre-running at 100 V for 20 minutes in TBE buffer. The low range ssRNA ladder from New England Biolabs was used as a reference in all gels, which were ran at 150 V for 45 minutes and subsequently stained with ethidium bromide.

### In vivo *toxicity assay*

*E. coli* BL21(DE3) cells transformed with *rnlA_pET28*, *R255A_rnlA_pET28*, *E258A_rnlA_pET28a*, *R318A_rnlA_pET28a, H323A_rnlA_pET28a* or *pET28a* (empty vector) were inoculated in 5 mL LB with kanamycin at a final concentration of 40 μg ml^−1^, at 310 K (200 rpm). After overnight incubation, cultures were diluted 100-fold in a honeycomb plate containing 297 μl LB medium, kanamycin (40 μg ml^−1^) and IPTG at 1 mM, (time point 0 in supplementary Figure S2). Growth curves were obtained by incubating the plates in a Bioscreen C Device at 310 K under continuous shaking. OD (595 nm) measurements were taken every 15 minutes for 12 h.

### Primer extension assays

0.08 μg of total MS2 RNA from each ribonuclease activity reaction mix were used for primer extension analysis with primer 5’-ttagtagatgccggagtttgctgcg-3’. The 5’-end of this primer is complementary to the stop codon of the MS2 coat protein mRNA, which is located between nucleotides 1338 and 1727. The primer was labeled on its 5’end with (γ ^32^P)-ATP (Perkin Elmer, 3000 Ci.mmol^−1^) and T4 polynucleotide kinase (Thermofisher). One pmol of labeled primer was mixed with one μl of ribonuclease reaction mix, after the reactions were carried out as described in the previous section except for the following modifications. 36 pmol of RnlA were incubated for 10 minutes with increasing amounts of RnlB, to evaluate the effect of the antitoxin on the RnlA specific cleavage of MS2 RNA. Additionally, other RnlA-MS2 reactions were incubated for 1, 2, 5, 10 and 20 minutes to monitor the kinetics of the enzymatic reaction. All reactions were stopped by the addition of a 2.5 molar excess of RnlB antitoxin. A range of concentrations for the RnlAΔ1-91 truncated protein, the interface-destabilizing RnlA mutants V206R and D245R, as well as 50 pmol of RnlA single-alanine mutant proteins were also included in primer extension analysis. These reactions were set up as described in the previous section, except that incubation of proteins and MS2 RNA was for 30 minutes. 36 and 50 pmol of wild-type RnlA, 100 pmol of RnlB only, and co-incubation of toxin and antitoxin prior to the addition of substrate were also included as references and negative controls. RnlAΔ1-91 and the V206R and D245R RnlA mutants preincubated with equimolar RnlB amounts were also included in the analysis.

Primer and template RNA mix were incubated for 5 minutes at 338 K and allowed to cool down to RT for approximately 30 minutes to favor annealing. Recombinant M-MuLV reverse transcriptase from the ProtoScript® II kit (NEB) was used to synthesize first strand cDNA from the annealed primer-MS2 RNA mix, according to the kit’s manufacturer. A set of sequence ladders for the MS2 coat protein was obtained by addition of the four ddNTPs (Sigma) in a final concentration of 1 mM each to the M-MulV reaction mix containing dNTPs at 0.25 mM each. Primer extension reactions were incubated at 315 K for one hour after which the enzyme was inactivated for 5 minutes at 353 K and subsequently stored at 253 K. Formamide dye mix was added to the samples and after incubation at 365 K for 3 minutes they were loaded on a 6 % denaturing polyacrylamide gel and ran for approximately 2 hours at 60 W (≈ 1500 V).

### ssDNA binding assays

The 5’-biotin-labeled oligonucleotide TTGCTGCGATTGCTGAGGGAATCGGGTTTCCATCTTT annealed to the 80-mer oligonucleotide of sequence <u>TTCCGACTGCGAGCTTATTGTTAAGGCAATGCAAGGTCTCCTA</u>AAAGATGGAAACCCGATTCCCTCAGCAATCGCAGCAA with free 3’-OH followed by a 43-nucleotides long ssDNA segment (underlined) which corresponds to the MS2 RNA sequence spanning over RnlA-specific cleavage sites 1 to 4 as identified in our primer extension assays. Biolayer interferometry (BLI) experiments were carried out in an Octet RED96 system with Streptavidin biosensors loaded with the primer-template DNA and blocked with BSA. The affinity of wild-type RnlA, the RnlAΔ1-91 truncate and single-alanine mutant proteins to this ssDNA oligonucleotide was measured by placing the sensors into solutions of several protein concentrations. PBS supplemented with 0.5 % BSA and 0.02 % Tween20 was used in all steps. Partial fitting of the data during association and dissociation to a 1:1 model (1 substrate for 1 RnlA dimer) was performed with the Octet Data Analysis software (FORTÉBIO), followed by the steady state analysis with the theoretical *R_eq_* values based on the curve fits. The analysis was performed in triplicate for each protein and the estimated *K_D_* values averaged and reported with associated errors.

### Macromolecular crystallography

Crystallization and data collection for His-RnlA and the RnlA-RnlB-His complex have been described (24). RnlA mutant R255A and the RnlAΔ1-91 truncate in 20 mM Tris-HCl pH 8, 150 mM NaCl, 1 mM TCEP were concentrated to 18 and 24 mg ml^−1^, respectively. Crystallization conditions were screened using a Mosquito HTS robot from TTP Labtech (http://ttplabtech.com/) using 0.1 μl of protein solution and 0.1 μl of reservoir solution in a sitting drop equilibrated against 100 μl of reservoir solution. Crystals of R255A RnlA grew after one week in 100 mM Tris-HCl, pH 8.5, 8 % PEG 8000. Crystals of RnlAΔ1-91 grew after 2-4 days in 0.2 M NaCl; 0.1 M BIS-Tris; pH 5.5; 25 % PEG 3350. Data were collected at beamline Proxima 2A of the Soleil Synchrotron (Gif-Sur-Yvette, France) for R255A RnlA mutant and Proxima 1 beamline of the Soleil synchrotron for RnlAΔ1-91. All data were analyzed using XDS (29) and are summarized in Supplementary Table S1. Additionally, merged intensity data for RnlAΔ1-91 were analyzed by the DEBYE and STARANISO software (30) via the web interface to perform Bayesian estimation of structure amplitudes, and to apply an anisotropic correction to the data. The estimation of the unit cell content was performed by the MATTHEW COEF program implemented in the CCP4 program suite (31).

The structure of the RnlAΔ1-91 truncation was determined by molecular replacement with Phaser. TBP2 and HEPN domains from PDB ID 4I80 were used as search models. Coordinates were refined in *phenix.refine* (33), using an intensities-based maximum likelihood target including rigid body and TLS refinement with automated group determination as implemented in *phenix.refine*. The structures of wild-type His-RnlA as well as of the R255A mutant protein were determined by molecular replacement with Phaser (32) using the deposited co-ordinates of the RnlA structure (PDB entry 4I8O) (7) determined in a different space group. NCS and refernce model (from the RnlAΔ1-91 structure) restraints were imposed in the refinement of RnlA WT and R255A structures. Rebuilding and addition of solvent atoms was done using Coot (34).

The structure of the RnlA:RnlB-His complex was determined by molecular replacement using Phaser-MR (35). A variety of search models was used ranging from the full RnlA dimer in PDB entry 4I8O to each of the individual domains of RnlA. Only the C-terminal RnlA domain (DBD, resid 203-357) resulted in an acceptable solution. Two copies of this domain were placed in a dimeric arrangement that did not reproduce the free RnlA dimer with LLG and TFZ values of 650 and 25.7, respectively. This partial solution did not provide sufficient phase information to allow building of the remaining parts of the structure. *phenix.mr_rosetta* (36) provided the essential tools for further model-building. The partial model and phases previously obtained from Phaser-MR were refined in Phenix and then used for a first phenix.mr_rosetta run. 9-mer and 3-mer fragments from the PDB, compatible with the sequence of RnlA were generated by the Robetta fragment server (http://robetta.bakerlab.org) and tallied as inputs to *phenix.mr_rosetta*.), Several cycles of phenix refinement and manual model building in Coot improved the phases and provided a new partial model, consisting now of 445 residues which was used for another cycle of *phenix.mr_rosetta*, this time also including the Robetta generated fragments for the RnlB sequence. The overall best solution of the Autobuild run of this *phenix.mr_rosetta* job rendered *R/R_free_* values of 0.29/0.32 with 570 residues built in. Further iterative cycles of refinement and both manual and automated model building using Coot and buccaneer (37), respectively, lead to the completion of the RnlA:RnlB structure. All refinement statistics are given in Supplementary Table S1.

### Small angle X-ray scattering

SEC-SAXS data were collected at beamlines BM29 (ESRF) and SWING (Soleil Synchrotron (Gif-Sur-Yvette, France)), using a setup with a SEC column (Shodex KW402.5-4F) placed just before the capillary to which the X-rays are directed. This setup helps to remove aggregates, that could interfere with the measurements, from the sample. Concentration series were collected in batch mode for RnlA wild-type and D245R mutant. Protein samples were prepared as described above for crystallization, concentrated and briefly centrifuged in a table-top centrifuge at 10 000 rpm for 2 mins, before loading on the SEC column or capillary. All samples were measured in 20 mM Tris, pH 8, 150 mM NaCl, 1 mM TCEP supplemented with 5 % glycerol for stabilization of proteins in solution, with a constant column flow of 0.2 ml min^−1^. The final scattering curve was generated for each sample in the SEC-SAXS runs after a range of scattering curves around the peak was normalized and averaged. The *R_g_* values were derived from the Guinier approximation at small *q* values while the *I_o_* parameter was estimated by extrapolation to *q = 0* using the ATSAS suite (38). For the samples measured in batch mode, 10 successive 1 second frames were collected for each concentration point in a series. The low angle data collected at lower concentration were extrapolated to infinite dilution and merged with the higher concentration data to yield the final scattering curves.

Computation of the SAXS profile of the atomic resolution models and its comparison to the experimental data was performed by FoXS webserver (39) (https://modbase.compbio.ucsf.edu/foxs/), which optimizes the excluded volume (*c_1_*) and hydration layer density (*c_2_*) to improve the fit of a model to the experimental SAXS data.

The MutiFoXS (39) tool of this webserver was used to address conformational heterogeneity in solution by considering multiple states contributing to the observed SAXS profile.

All-atoms input models were generated by modeling missing fragments (N-terminal histidine tags and missing loops) with MODELLER (https://salilab.org/modeller/) (40). 10000 conformations were generated by sampling the space of the φ and ψ main chain dihedral angles of selected flexible residues with a Rapidly exploring Random Trees (RRTs) algorithm. Residues in loops and connecting domains were defined as flexible while keeping the rest of the protein as rigid body. Subsequently, the computation of the SAXS profile for each sampled conformation was performed followed by the enumeration of the best-scoring multi-state models using the multi-state scoring function implemented in MultiFoXS (39). SAXS data collection and processing, as well as modeling parameters are summarized in Supplementary Tables S2-S5.

## RESULTS

### RnlA is a HEPN protein that adopts a non-canonical dimer in its resting state

Expression of a bicistronic construct encoding N-terminally His-tagged RnlA (His-RnlA) followed by RnlB allows the purification and crystallization of His-RnlA, since RnlB is largely lost during purification possibly due to proteolysis. The resulting structure determined at 2.9 Å is very similar, although in a different crystal form, to an RnlA structure (PDB ID 4I8O) presented earlier (7) (RMSD = 0.907 Å over 4418 atoms from 668 residues). The RnlA monomer consists of three domains, previously denoted N-terminal domain (NTD), N-terminal repeated domain (NRD) and Dmd-antitoxin binding domain (DBD) (residues 1-92, 93-193, and 194-357, respectively, Figure 1a). A DALI search revealed that the NTD and NRD domains adopt a TATA box-binding protein (TBP)-like domain fold most similar to the N-terminal domain of *Aquifex aeolicus* RNase H3 (41) (PDB ID 3VN5, Z score of 6.2 - Supplementary Figure S1a). Therefore, they will be referred to as TBP1 and TBP2 from hereon. The C-terminal domain adopts a HEPN nuclease fold and will be referred to as the HEPN domain. The closest structural relatives for the HEPN domain, aside from LsoA, comprise the TA toxins SO_3166 and HI0074 from *Shewanella oneidensis* and *Haemophilus influenza,* respectively (21,42), but also other HEPN proteins including human Sacsin are readily identified in a DALI search (43).

**Figure 1:**
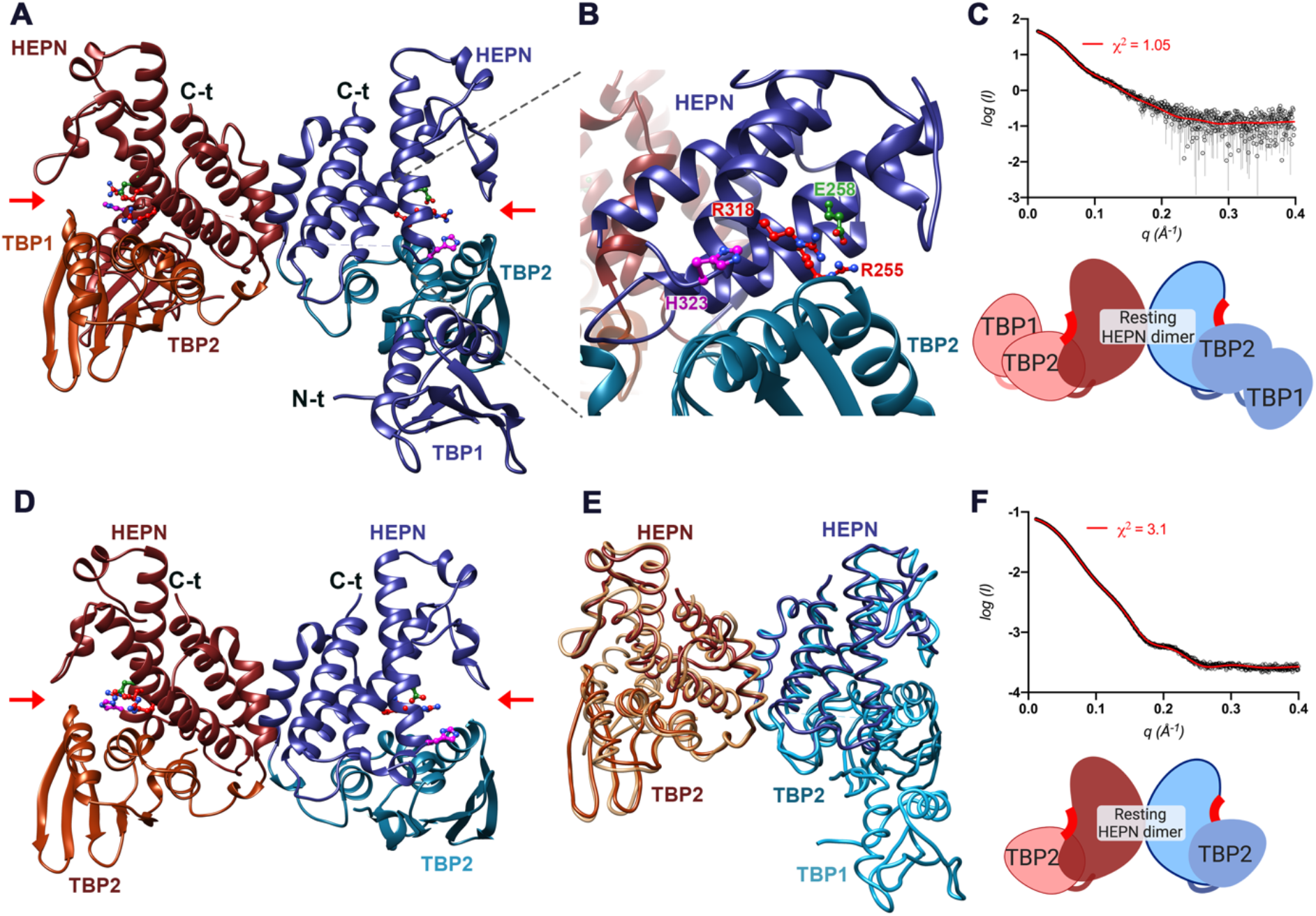
Unbound RnlA exists as a resting dimer both in the crystal and in solution. **A** Ribbon representation of the RnlA dimer in its free state. The three domains are labeled in both chains (TBP1, TBP2, HEPN). Chains are colored in red and blue; TBP2 domains are colored in a different shade of red and blue, respectively, to mark a clear distinction between domains. Red arrows indicate the position of the conserved catalytic residues in the TBP2-HEPN interface of each chain. **B** Ball and stick representation of the proposed catalytic residues R255, E258, R318 and H323 in the TBP2-HEPN interface. **C, F** Measured SAXS profiles (empty circles) for the isolated RnlA and RnlAΔ1-91 proteins with the theoretical profiles (red line) calculated for all-atom models of their respective crystallographic structures. *χ*^2^ values for the comparisons between experimental and calculated SAXS profiles are indicated. Cartoon representations of the domain organization of RnlA full-length and the truncated RnlAΔ1-91 version are included below the curves. **D** Ribbon representation of the crystallographic structure of the RnlAΔ1-91 truncated variant. Coloring is as in panel A. **E** Trace representation of the superposition of full-length RnlA dimer and the RnlAΔ1-91 truncated variant. RMSD between both structures is 1.56 Å over 435 Cα.

A sequence alignment of the RnlA family (PF19034) in the *pfam* database (http://pfam.xfam.org) reveals the presence of an RX_4-6_H motif (R318 and H323 in RnlA) that was earlier identified among members of the HEPN superfamily and that is considered to be part of the active site of HEPN ribonucleases. This minimal motif is located at the C-terminus of helix α5 of the HEPN domain (α13 in full length RnlA) but differs in length in the different HEPN subfamilies (16–18). In addition, an arginine and glutamate residue (R255 and E258) that were suggested as possible catalytic residues in LsoA (8) are equally conserved in the RnlA/LsoA subfamily, but not in other HEPN subfamilies (Supplementary Figure S1b).

In this structure, RnlA dimerizes via its HEPN domain, but a non-canonical HEPN dimer is formed via interactions between helixes α8 and α10. We will refer to this conformation as the resting or unbound state as it is not involved in binding of substrate nor antitoxin (see later). The presence of this non-canonical dimer is confirmed in solution by SAXS (Figure 1a,c), and is in agreement with previous results (7). Although the dimer interface is relatively small for a protein of this molecular weight (1638 Å^2^ buried), PISA analysis of the full-length RnlA structure outputs this dimer conformation as the only stable assembly among the different contacts in the crystal, with a theoretical free Gibson’s energy gain of −15 kcal mol^−1^.

Further evidence for the stability of this non-canonical HEPN dimer comes from a truncation mutant of RnlA (RnlAΔ1-91) that lacks the N-terminal residues 1-91 corresponding to the TBP1 domain. The crystal structure of RnlAΔ1-91 shows the same non-canonical HEPN dimer (Figure 1d,e - RMSD of 1.56 Å over 435 common Cα atoms with the full-length dimer) and was confirmed in solution using SAXS. The theoretical SAXS profile for an all-atom model of the RnlAΔ1-91 crystallographic dimer agrees with the experimental SAXS profile with *χ*^2^ = 3.1 (Figure 1f).

### RnlA is a ribonuclease with broad sequence specificity

To characterize the endoribonuclease activity of RnlA, the 3569 nucleotides-long RNA from bacteriophage MS2 was used in an *in vitro* RNase assay. Wild-type RnlA efficiently degrades this substrate, revealing multiple bands on a denaturing gel ranging from 500 to less than 50 nucleotides in size (Supplementary Figure S2a). Moreover, RnlA is fully inhibited by the addition of a twofold excess of RnlB prior to the addition of MS2 substrate. RnlB on its own does not show any ribonuclease activity under the same conditions.

The specificity of wild-type RnlA was investigated by primer extension with a 5’-^32^P-labeled primer annealing to the coat protein mRNA on MS2 genome. Compared to control reactions lacking RnlA, several RnlA-cleavage specific bands are observed from 1 minute of incubation with RnlA onwards. The intensities of the higher molecular weight RnlA-specific bands decrease as the incubation time increases, which indicates the subsequent degradation of RNA fragments by the endoribonuclease activity of RnlA (Figure 2a). When RnlA is incubated with increasing amounts of RnlB prior to the addition of MS2 RNA substrate, the RnlA-specific cleavage bands disappear or show background intensity (Figure 2b).

**Figure 2:**
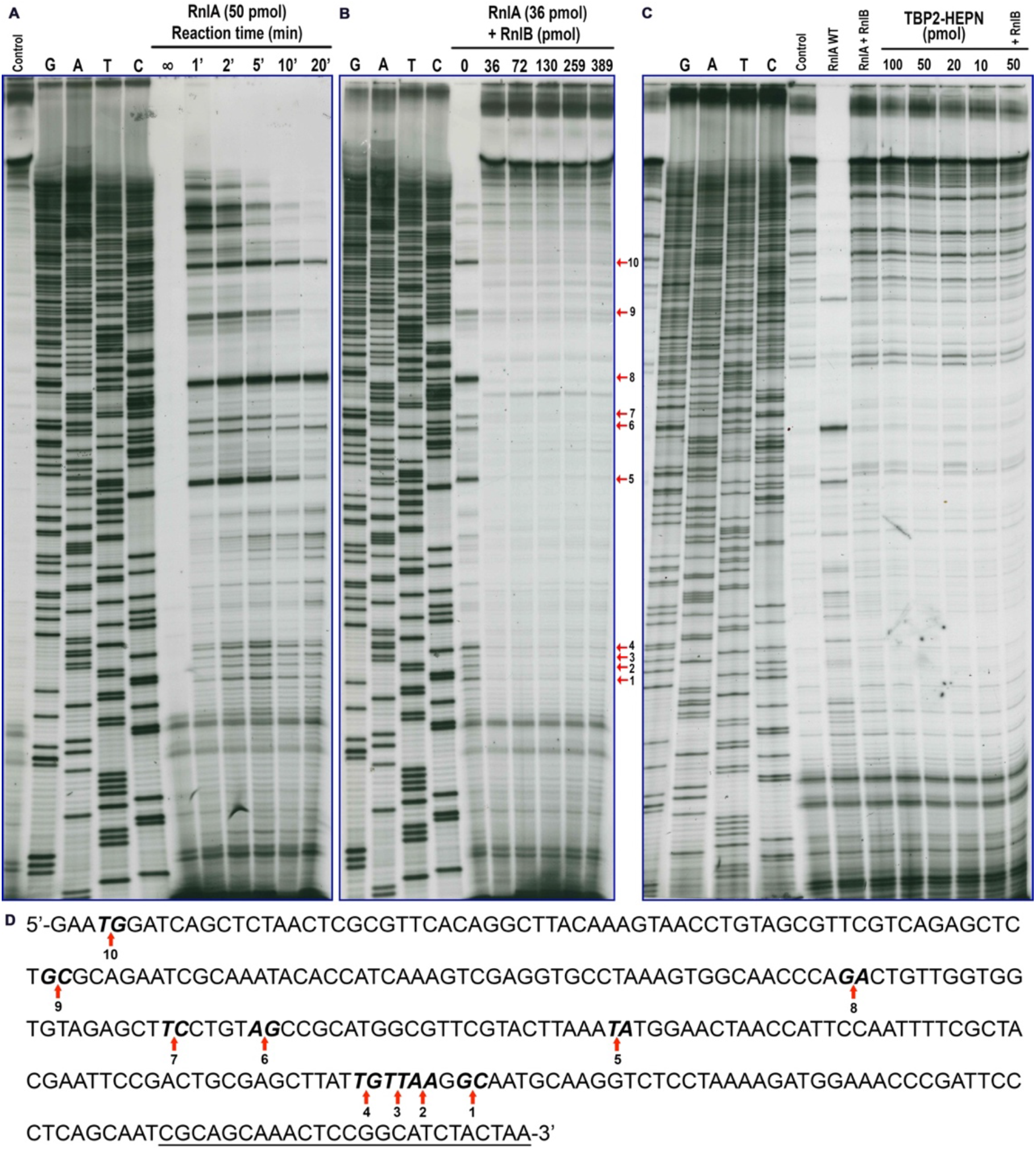
Specificity of RnlA ribonuclease activity. Autoradiographs of the coat-protein MS2-RNA primer extension analysis. **A** RnlA activity time-progress. Purified His-RnlA was incubated with MS2 RNA substrate for increasing time periods prior to primer extension analysis with 5’-^32^P-labeled primer. All reactions were stopped by a 2.5 molar excess of RnlB antitoxin, except for one marked as infinite time. **B** Effect of RnlB on the endoribonuclease activity of RnlA. Increasing amounts of RnlB were incubated with RnlA for 10 minutes prior to the addition of MS2 substrate. Subsequent primer extension reactions were carried out as in a. RnlA-specific cleavage bands are marked by arrows and numbered. **C** Decreasing amounts of RnlAΔ1-91 truncated protein was incubated with MS2 RNA substrate for 30 minutes prior to primer extension analysis performed as in a. Control reactions consisting of no protein, RnlA, RnlA preincubated with RnlB antitoxin and RnlAΔ1-91 preincubated with RnlB were included for comparison in the same primer extension analysis. All reactions were carried out in parallel. Sequence ladders were obtained by the addition of the four ddNTPs to primer extension reactions. **D** cDNA sequence annotation of coat-protein region on MS2 RNA. The 10 RnlA-specific cleavage bands identified in B are marked by arrows and bases are in bold. The 3’ under-scripted end represents the labeled primer.

Ten strong RnlA cleavage sites can be identified in the primer extension analysis. They are indicated by numbered arrows on the sequencing gel and corresponding inter-nucleotide positions on the MS2 cDNA sequence fragment (Figure 2b,d). Most but not all of the cleavage sites occur within or at the ends of base-paired RNA stretches, according to the predicted secondary structure of this region of the MS2 RNA performed in the RNAfold WebServer (http://rna.tbi.univie.ac.at/). Cleavage sites 1 to 4 are remarkably close on the sequence, with cleavages 2 to 4 occurring every second nucleotide. This region of the MS2 RNA might be particularly prone to RnlA cleavage. In terms of sequence specificity, no consensus could be generated by simple sequence inspection. RnlA seems to be able to cleave both downstream of purines and pyrimidines, which agrees with the broad sequence specificity previously suggested (5). Based on these data, it cannot be excluded that substrate specificity of RnlA is either determined by RNA structure rather than sequence, or that RnlA is an exonuclease where the strongest bands in the primer extension analysis correspond to pauses due to stable structure in the substrate.

### The N-terminal domain of RnlA is required for substrate binding

To assess the function of the N-terminal TBP1 domain in the ribonuclease activity of RnlA, primer extension experiments were performed with the RnlAΔ1-91 truncate lacking this domain. Surprisingly, no activity could be detected at concentrations varying from 10 to 100 pmol, nor did the addition of RnlB antitoxin influence the outcome (Figure 2c). These results are unexpected, considering the presence of a supposedly intact catalytic site in the RnlAΔ1-91 variant.

Therefore, substrate-binding experiments were performed with the RnlAΔ1-91 truncation variant. We observe a two-order decrease in binding affinity when the TBP1 domain is absent compared to the full-length protein (Table 1, Supplementary Figure S3). These findings indicate that the TBP1 is required for substrate binding and explain why no activity is observed when RnlAΔ1-91 is used. They also provide a potential explanation for the seeming lack of RNA specificity of RnlA, namely that RNA sequence specificity is at least in part embedded in the TBP1 domain, but that the sequence that is recognized as such is too far removed from where the HEPN domain actually cuts to be readily identified.

**Table 1:** TBP1 is required for substrate binding and Arg255 is involved in substrate binding. Binding of RnlA and its single alanine mutants to ssDNA. Data were collected in an Octet RED96 instrument and analyzed with the manufacturer data analysis tool v. 9.0 (FORTÉBIO). K_D_ values are derived from the steady state analysis of the data fits to a 1:1 model and R_Max_ values are extrapolated from the fits to this model.

| Protein | $K_D$ (M) |
| --- | --- |
| RnIA | $4.30 \cdot 10^{-6} \pm 3.20 \cdot 10^{-6}$ |
| R255A RnIA | $1.97 \cdot 10^{-4} \pm 1.13 \cdot 10^{-4}$ |
| E258A RnIA | $7.17 \cdot 10^{-6} \pm 1.91 \cdot 10^{-6}$ |
| R318A RnIA | $6.93 \cdot 10^{-6} \pm 1.37 \cdot 10^{-6}$ |
| RnIAΔ1-91 | $4.83 \cdot 10^{-4} \pm 3.29 \cdot 10^{-4}$ |

### R255, E258, R318 and H323 are essential for activity

Single-alanine substitution mutants of the four putative catalytic residues of RnlA suggested above do not show any activity in the presence of MS2 RNA under the same conditions used for wild-type RnlA (Supplementary Figure S2a). Together with the high conservation scores for these residues in the RnlA/LsoA family and their involvement in the toxicity of LsoA *in vivo* (8), this finding constitutes a strong indication of a role for R255, E258, R318 and H323 in catalysis or substrate binding.

Additionally, these mutants were included in the primer extension analysis to further confirm the lack of activity observed in the previous MS2 RNA cleavage assay. As seen on Figure 3a, cDNA amplification of MS2 RNA after incubation with each of the single-alanine RnlA mutant proteins shows the same background intensity pattern as negative controls in absence or RnlA or presence of RnlB incubation. This result confirms the absence of endoribonuclease activity for these RnlA mutants and of RnlB.

**Figure 3:**
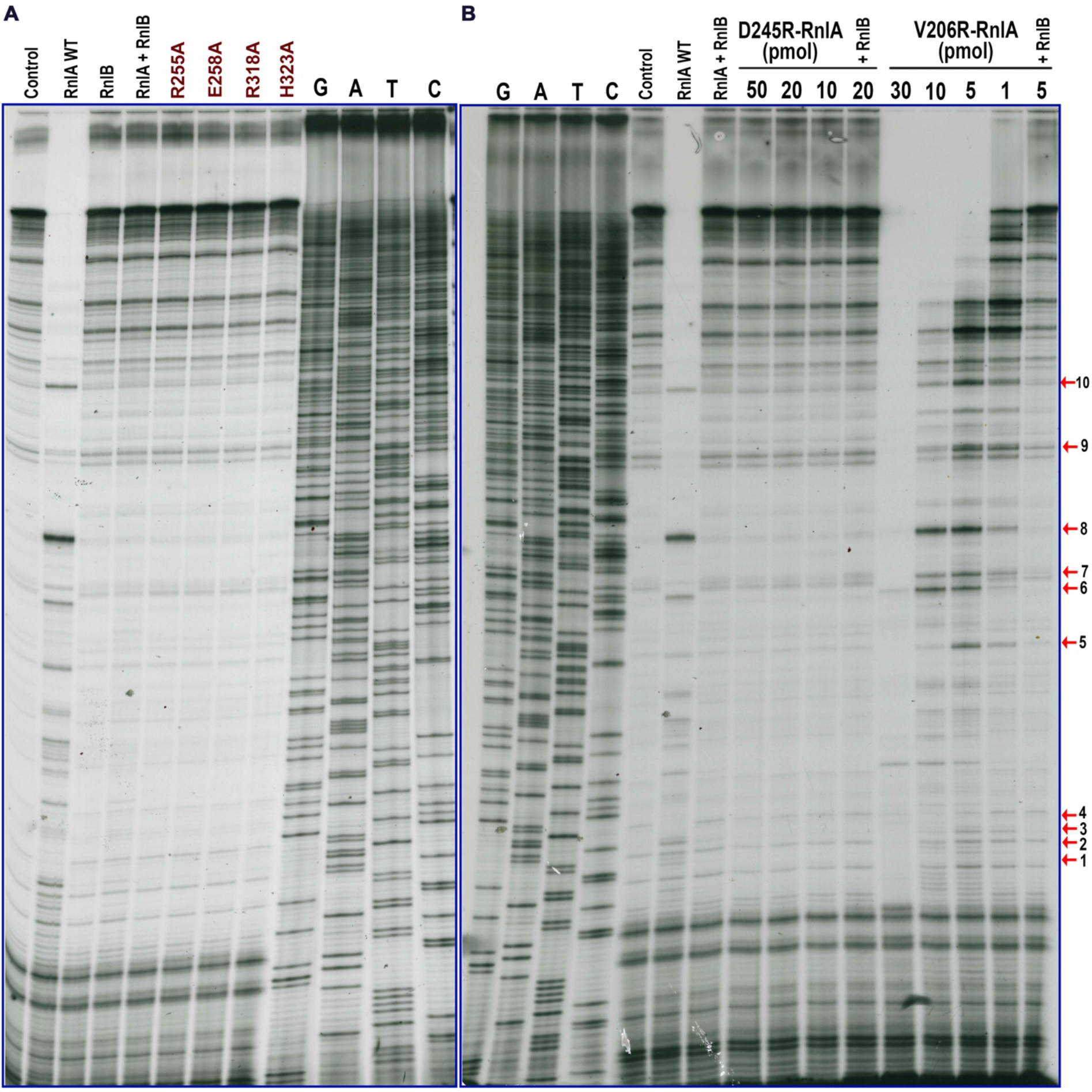
HEPN conserved motif defines the catalytic site of RnlA and canonical dimerization is required for activity. Autoradiographs of the coat-protein MS2-RNA primer extension analysis. **A** R255A, E258A, R318A and H323A RnlA mutants were incubated with MS2 RNA substrate for 30 minutes prior to primer extension analysis performed as in Figure 2. **B** Decreasing amounts of interface-destabilizing mutants D245R and V236R RnlA were incubated with MS2 RNA prior to primer extension as in a. V206R and D245R RnlA mutants preincubated with equimolar RnlB amounts were also included in the analysis. RnlA-specific cleavage bands are marked by arrows and numbered. Control reactions consisting of no protein, RnlA wild-type, RnlB antitoxin and RnlA:RnlB complex were included for comparison in the same primer extension analysis. All reactions were carried out in parallel. Sequence ladders were obtained by the addition of the four ddNTPs to primer extension reactions.

To validate our single alanine mutants *in vivo*, wild-type and mutants were expressed from *pET28a* in *E. coli* BL21(DE3). We observe an increase in lag time for growth of the cells over-expressing wild-type RnlA compared to the empty vector control. This result is in agreement with earlier measurements that indicate that RnlA slows down but does not prevent bacterial growth (5). No such growth delay is observed for R255A, E258A, R318A or H323A RnlA expressing cells (Supplementary Figure S2b).

In order to explore the role of these four highly conserved residues in substrate binding, a 43-nucleotides ssDNA segment comprising RnlA-specific cleavage sites 1 to 4 on the MS2 RNA sequence was used in Biolayer Interferometry (BLI) experiments (Supplementary Figure S3). While the differences in conformational preferences between DNA and RNA will certainly lead to significant differences in affinity, the use of ssDNA has the advantage that cleavage is fully prevented. This approach is valid as only the protein and not the substrate is varied in this experiment and we are interested in relative rather than absolute affinities. A similar approach was earlier adopted to study substrate recognition of *Staphylococcus aureus* MazF (28).

Wild-type RnlA binds to this substrate mimic with a *K_D_* in the μM range. For the mutants E258A and R318A, this affinity does not drastically change, indicating that these two residues are not crucial for substrate binding but may have a catalytic role (Table 1 and Supplementary Figure S3). On the other hand, for mutant R255A the affinity for the substrate mimic is weakened by two orders of magnitude, which points to the involvement of this arginine residue in substrate binding (Table 1 and Supplementary Figure S3a,b). BLI analysis with mutant H323A was not possible due to protein instability at the concentrations required for the assay.

### RnlB antitoxin binds a canonical HEPN dimer of RnlA

To understand how RnlB antitoxin inhibits RnlA, we determined the crystal structure of the RnlA:RnlB complex at 2.6 Å resolution. This structure shows one RnlA dimer interacting with two monomeric RnlB molecules (Figure 4a). Furthermore, this hetero-tetramer is representative of the RnlA:RnlB assembly in solution as confirmed by SAXS (Supplementary Figure S4). RnlB is entirely folded and consists of a four-stranded mixed β-sheet sandwiched between two α-helices (Figure 4, Supplementary Figure S5a). The most striking feature of the RnlA:RnlB complex is that here RnlA forms a dimer that is fundamentally different from what is seen in the free RnlA crystal structure. The dimer is again formed via the HEPN domain but resembles the V-shaped canonical dimer that has been observed in different HEPN subfamilies. This novel arrangement brings R255, E258, R318 and H323 from helix *α*13 in both monomers together, with R318 and H323 remaining exposed but R255 becoming buried and mediating inter-monomer contacts. In this arrangement, the role of E258 seems to help to correctly position the side chain of H323.

**Figure 4:**
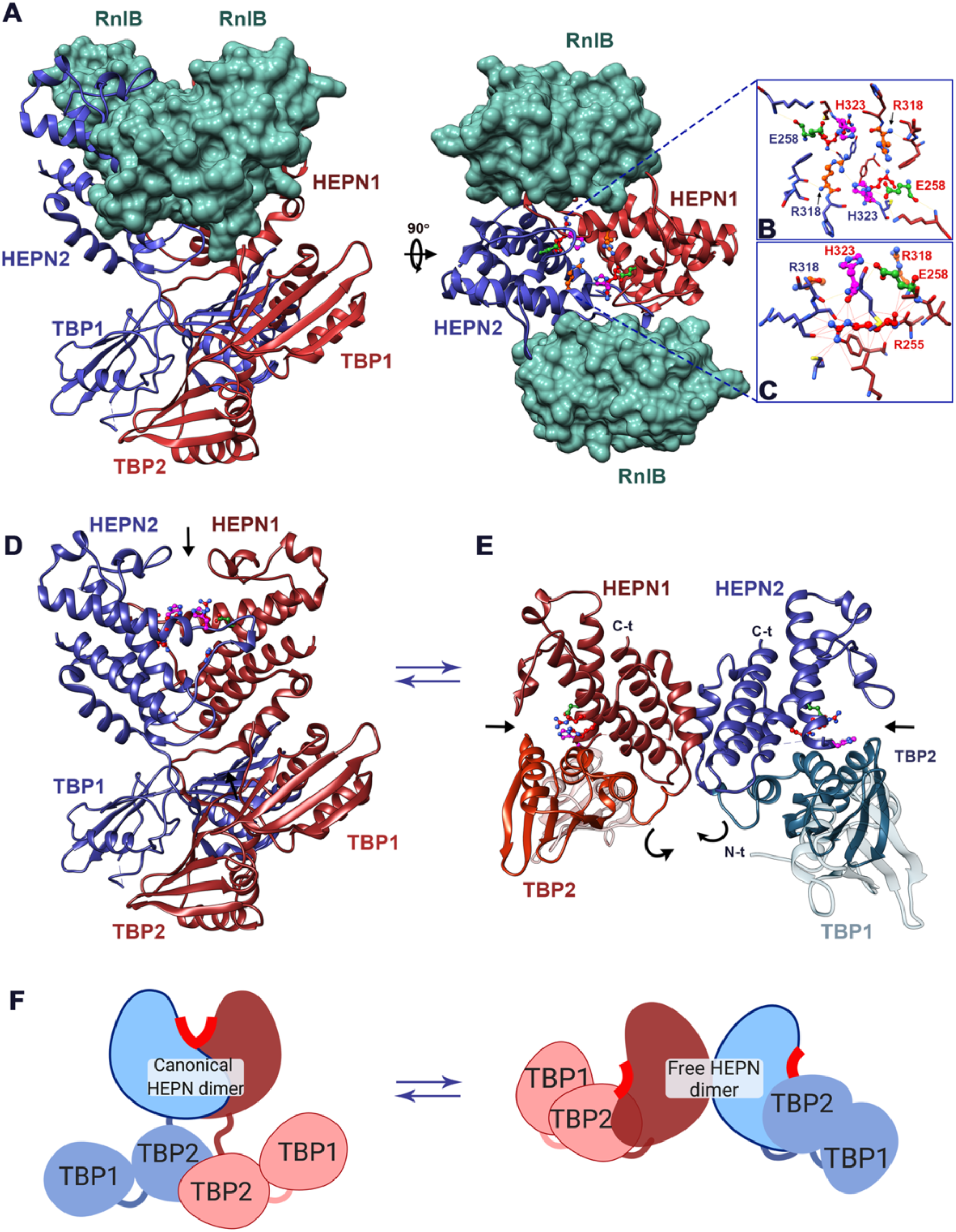
RnlB recognizes a canonical HEPN RnlA dimer. **A** Ribbon representation of the RnlA:RnlB heterotetrameric complex. Chains in the RnlA dimer are colored in red and blue, while the two RnlB monomers are shown in green surface representation. The three domains (TBP1, TBP2, HEPN) are indicated. Top view of the RnlA:RnlB complex. **B** Catalytic residues are shown in ball and stick and colored by heteroatom. Relevant contact residues are represented as sticks. **C** Contact residues of buried R255 (in chain A) are represented as sticks and colored by heteroatom. Contacts (Van der Waals radii overlap > 0.4 Å) are represented as red lines. Labels are displayed and H-bonds are shown as yellow lines. Ribbon representation of RnlA dimers in the RnlB-bound (**D**) and free (**E**) forms. Coloring is the same as in panel A. Black arrows indicate the position of the catalytic residues in each dimer. Catalytic residues are represented as ball and stick and colored by heteroatom. **F** Cartoon representation of the domain organization and dimerization modes of antitoxin-induced RnlA dimer (canonical HEPN dimer) and free RnlA dimer (free HEPN dimer).

The new contact interface formed between the HEPN domains results in the burial of 4070 Å^2^, which is significantly higher than the 1638 Å^2^ of the HEPN dimerization interface in the unbound state of RnlA (Figure 4b,c). At the same time, the TBP2 domains detach from the HEPN domains and also associate into a domain-swapped dimeric arrangement that buries another 2776 Å^2^. This results in a large 12-stranded cross-domain β-sheet (Supplementary Figure S6). Finally, each RnlB further stabilizes the novel arrangement by bridging the HEPN dimer and via its interactions orders the otherwise disordered loop T326-D336 of RnlA.

### The canonical HEPN dimer is the active conformation of RnlA

In order to determine whether canonical HEPN dimerization is required for activity, mutations in the HEPN domain were designed to destabilize the resting or the canonical HEPN dimer conformations. V206R RnlA was designed to interfere with the formation of the resting state dimer. Importantly, this residue is exposed to the solvent in the HEPN canonical dimer but sterically prevents the formation of the resting dimer. On the other hand, D245R RnlA destabilizes the canonical HEPN dimer in the antitoxin-induced RnlA conformation but should not interfere with the formation of the resting RnlA dimer. Additionally, the already mentioned R255A mutation would remove interactions at the canonical HEPN dimer interface and may thus stabilize the resting dimer conformation. SAXS data for both D245R and R255A RnlA as well as the crystal structure of R255A RnlA show that these mutants indeed adopt the resting RnlA dimer conformation (Figure 5, Supplementary Figure S7). The V206R mutant unfortunately is too insoluble to determine its dominant solution state by SAXS.

**Figure 5:**
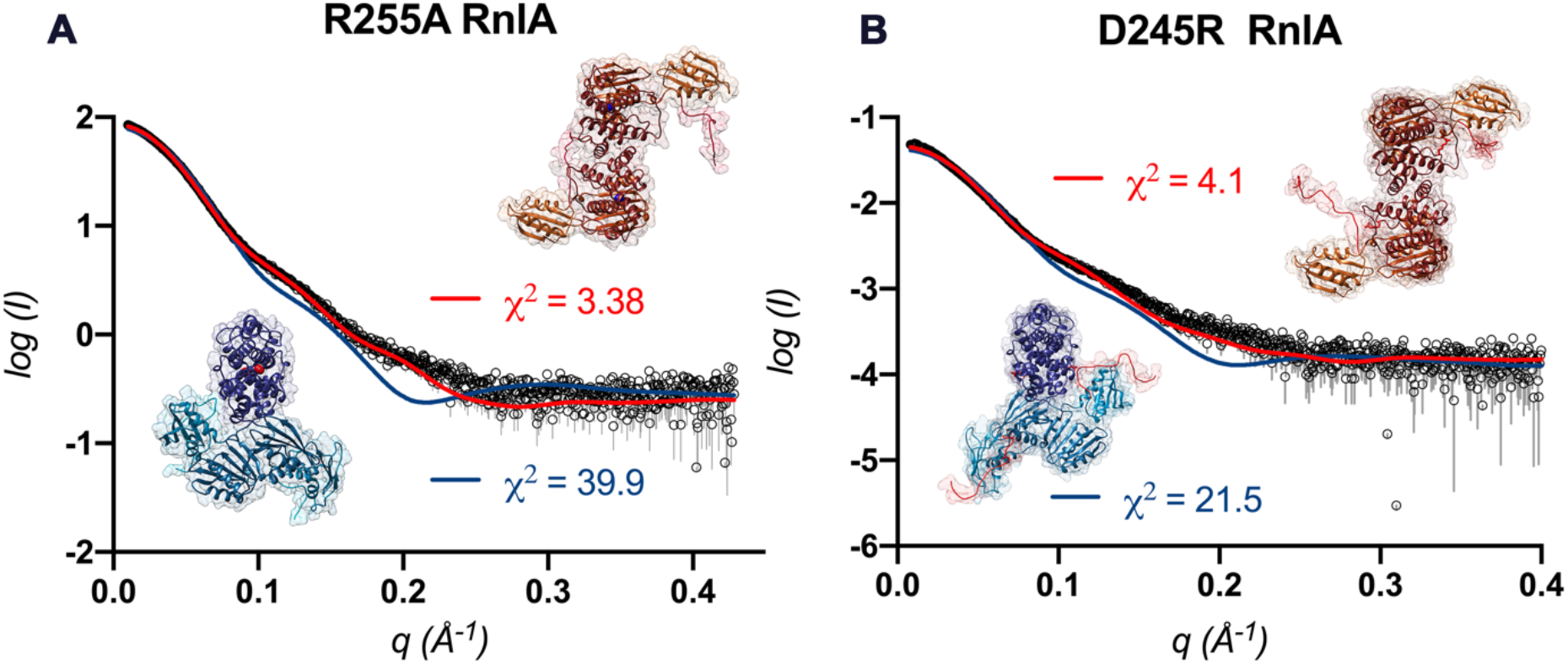
Mutations in residues involved in dimerization shift the equilibrium in solution between the free and antitoxin-induced RnlA conformations. Small angle X-Ray Scattering (SAXS) measurements of R255A RnlA (**A**) and D245R RnlA **(B**). All panels show the comparison of the experimental data (open circles) to the theoretical profile for the corresponding all-atoms model obtained by the program MODELLER. MultiFoXS-calculated scattering curves for free RnlA (red curves) and RnlB-induced RnlA dimer (blue curves) conformations are compared to the experimental data for R255A and D245R RnlA. *χ*^2^ values for comparisons to free RnlA models are shown in red and to RnlB-induced RnlA models are shown in blue.

*In vitro* ribonuclease activity and primer extension assays were performed with these mutant proteins. V206R RnlA displays the same RnlA wild-type cleavage pattern in the primer extension experiments. Importantly, its activity is inhibited by the addition of RnlB antitoxin (Figure 3b), showing that it indeed can adopt the canonical HEPN dimer conformation. On the other hand, ribonuclease activity of D245R RnlA could not be detected at any concentration within a range of 10 to 50 pmol of protein (Figure 3b). As expected, the addition of RnlB antitoxin to this mutant protein did not cause a change in its activity. We next tested these mutants in our *in vivo* growth assay. In agreement with these results, the V206R mutant is active *in vivo*, inducing the same growth defect as the wild-type protein. In contrast, the D245R shows no active phenotype and growth of this mutant follows that of the empty vector control (Supplementary Figure S2c).

Taken together, these results indicate that HEPN canonical dimerization is required for activity, in contradiction to what had been previously reported for RnlA (7). HEPN canonical dimerization leads to the formation of a composite active site generated by the juxtaposition of the highly conserved residues located within a catalytic cleft (Supplementary Figure S8b).

## DISCUSSION

The HEPN domain is widespread across the proteomes of bacteria, archaea and higher eukaryotes. It shows a conserved topology despite high sequence divergence. HEPN domains are RNases that are characterized by a conserved RX_4-6_H active site motif. They are found in bacterial anticodon nucleases (ACNases) RloC (44) and PrrC (45) involved in the restriction of T4 phage, in the kinase-extension nuclease (KEN) domain of RNase L in vertebrates (16), in domains in the abortive infection genes (Abi) system and in domains encoded by genes related to the CRISPR_Cas loci (6). Several subfamilies of HEPN domains have been identified based on their structural variability outside the common HEPN core. RnlA and LsoA represent one of these subfamilies.

HEPN domains are dimerization units. From the crystal structures of different HEPN domain containing proteins, a canonical dimer association emerges. This association results in a V-shaped structure that brings together the RX_4-6_H motifs of both chains and forms a positively charged surface to which the RNA substrate docks. The details of this association vary between subfamilies, resulting in a range of more open or closed architectures in addition to a few cases that deviate from this canonical arrangement (Supplementary Figure S9). RnlA, although being a HEPN ribonuclease, was believed not to adopt the canonical HEPN dimer. The crystal structure of RnlA in its unbound state shows an alternative non-canonical HEPN domain-mediated dimer (7) while the homologous LsoA in complex with Dmd does not dimerize (8). In these structures the RX_4-6_H motif as well as R255 and E258 (or their equivalents in LsoA) are located on the interface between the HEPN domain and the TBP2 domain (Figure 1a,b; Figure 4c and Supplementary Figure S8a). As the equivalents of R255, E258 and R318 in LsoA are essential for its activity *in vivo* and the inhibitor Dmd locates in the groove between the HEPN and the TBP2 domains of LsoA, it was assumed that this corresponded to an active conformation of the enzyme where Dmd inhibits the protein by blocking the active site (8) (Supplementary Figure S10b,c). We now observe that RnlA when bound to its cognate antitoxin RnlB adopts a second HEPN-mediated dimer that resembles a canonical HEPN dimer, which has been proposed to represent the active conformation of other HEPN ribonucleases (Figure 4a,b and Supplementary movie 1). This dimer brings together the RX_4-6_H motifs from both monomers, as is the case for HEPN ribonucleases in general (17, 21) (Figure 4a,b and Supplementary Figures. S5c, S9). Thus, the change in quaternary structure between the unbound and RnlB-bound form of RnlA is drastic and to our knowledge has not been observed in any other protein. Not only does the dimer interface completely change with residues belonging to opposite sides of the HEPN domain monomer being involved, it also coincides with further domain rearrangements: the TBP2 domain detaches from the HEPN domain (thus making available its alternative dimer interface) and also dimerizes via swapping of its N-terminal β-strand (Supplementary Figure S6). Stabilization of the canonical HEPN dimer conformation leads to an active enzyme both in *vivo* and *in vitro*. In this arrangement, R255, which is not conserved among the HEPN superfamily, would not be a catalytic residue as suggested earlier (7), but contribute to stabilization of the canonical HEPN dimer (Figure 4a,b and Supplementary Figure S8b), in agreement with the observed defect in substrate binding of the R255A mutant.

RnlB binds to this active conformation of RnlA. Yet, RnlB does not bind to the active site groove of the HEPN domain. Rather, it is positioned adjacent to it, seemingly blocking entrance and exit of the symmetric substrate-binding groove (Supplementary Figure S5c). The side chains of E258, R318 and H323 remain at least partially exposed upon RnlB binding. This contrasts with SO_3166 from *Shewanella oneidensis*, the only HEPN protein in a toxin-antitoxin context for which a crystal structure is available, besides RnlA and its homolog LsoA (21). SO_3166 together with SO_3165 constitute a type II HEPN-MNT TA system. SO_3166 consists of a single HEPN domain that adopts the canonical HEPN-dimer arrangement, similar to the conformation in the RnlB-inhibited form of RnlA (Supplementary Figure S9). The presumed active site groove formed upon SO_3166 homodimerization is occupied by α-helix 4 of the antitoxin SO_3165, which obviously results in inhibition as the RX_4-6_H motifs are inaccessible to the substrate.

Although the canonical HEPN dimer of RnlA has a much larger interface than the alternative non-canonical resting dimer, the latter is the dominant species in solution in absence of any ligand as evidenced by SAXS. The stability of this “resting” dimer likely comes from interactions between the HEPN and TBP2 domains, which need to be broken before the canonical HEPN dimer can be formed. Using site-specific mutants that stabilize one of the two possible conformations, we could show that RnlA is active only when it forms the canonical HEPN dimer. A mutant that stabilizes the resting state by preventing canonical HEPN dimerization is inactive. Whether the alternative resting dimer, that exists in the crystal and in solution, has any function *in vivo*, e.g. by reducing activity and thus toxicity of the protein, remains unclear. In this context, it is relevant to note that RnlA and its relative LsoA are quite mild toxins compared to other RNase toxins, as they only slow down growth in absence of their antitoxin. Finally, our data for the first time point towards a functional role of the N-terminal TBP1 domain of RnlA. Deletion of this domain leads to an inactive enzyme that is defective in substrate binding. Possibly, this domain provides an anchor point on the larger RNA substrate, bringing the catalytic HEPN domain close to its substrate. As any specific recognition sequence for the TBP1 domain would be relatively far from the actual cutting site(s), this could explain the apparent lack of sequence specificity of RnlA. In agreement with such a function is that TBP domains are found associated with other RNases as well. In particular, the N-terminal domain of RNase H3, which is otherwise unrelated to RnlA, is such a domain. In the crystal structure of *Thermovibrio ammonificans* RNase H3 (46) (PDB ID 4py5), which is the only one in complex with a substrate, the 15 bp hybrid DNA/RNA duplex is sandwiched between the TBP domain and the RNase H3 catalytic domain. This TBP domain thus seems to have a role in substrate positioning, strengthening our assumption for a similar role in RnlA and LsoA.

## CONCLUSION

In contrast to what was concluded in earlier studies, RnlA is a bona-fide HEPN ribonuclease that requires canonical HEPN dimerization for activity. Drastic conformational changes mediate transitions between the active HEPN canonical form and the resting state that is adopted as the major conformation in solution when no RnlB or substrate is bound. This resting state may help to tune down its toxicity *in vivo*. The N-terminal TBP1 domain is required for substrate binding while interactions between the TBP2 and HEPN domains shift the conformational equilibrium of the protein towards the resting state. The antitoxin RnlB binds RnlA in its active state but prevents substrate binding via blocking the entrance and exit of the substrate binding groove (Figure 6). This study therefore reconciles the mechanism and regulation of RnlA with that of other HEPN ribonucleases.

**Figure 6:**
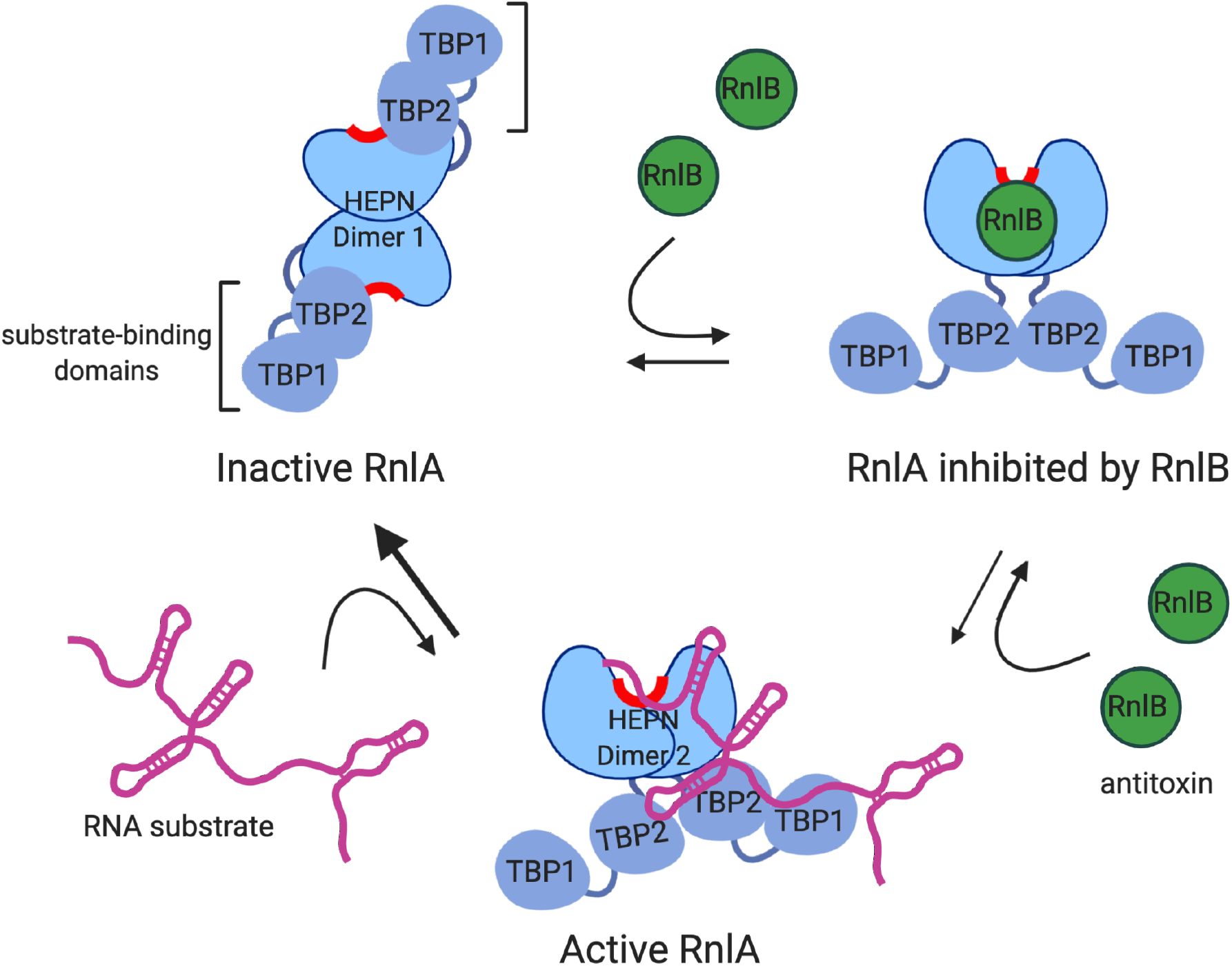
Model for the activity and inhibition of HEPN ribonuclease RnlA. RnlA drastically shifts domain orientation and dimerization interface between different states. RnlA free resting state involves the formation of HEPN dimer 1, which positions the conserved catalytic residues in the TBP2-HEPN interface of each chain in the dimer. The TBP1 and TBP2 adopt a TBP-like domain fold which is involved in substrate binding in other RNases. The TBP1 of RnlA is required for substrate binding and activity. The catalytically active RnlA conformation requires HEPN canonical dimerization (HEPN Dimer 2), which involves juxtaposition of the conserved catalytic residues in the formation of a composite catalytic cleft. The TBP1 and TBP2 assist in substrate binding and potentially in positioning the RNA substrate for catalysis. RnlB antitoxin recognizes the HEPN canonical dimer and binds on either side of the catalytic cleft, blocking substrate access. RnlB shifts the equilibrium of the enzyme from the free state to the RnlA canonical HEPN dimer.

## ACCESSION NUMBERS

All crystallographic data have been deposited into the Protein Data Bank (www.rcsb.org) The accession codes are the following: RnlA:RnlB, 6Y2P; RnlA, 6Y2Q; R255A RnlA, 6Y2R, RnlAΔ1-91, 7AEX. The solution scattering data has equally been deposited at the www.sasbdb.org database with the following accession codes: RnlA:RnlB, SASDHW7; RnlA, SASDHX7; RnlAΔ1-91 (TBP2-HEPN), SASDKL2; R255A RnlA, SASDHY7; D245R RnlA, SASDKS9.

## SUPPLEMENTARY DATA

Supplementary Data are available online.

## FUNDING

This work was supported by THE FWO-Vlaanderen [grant G.0B25.15N]; and the Onderzoeksraad of the Vrije Universiteit Brussel [SRP13].

## ACKNOWLEDGEMENTS

The authors thank Sarah Haesaerts for technical assistance and the beamline staff of the Soleil macromolecular crystallography beamlines for help during data collection and processing.

## CONFLICT OF INTEREST

The authors have no competing interests to declare.

## Notes

### Competing Interest Statement

The authors have declared no competing interest.

