## Supplementary Data for "Alternative dimerization is required for activity and inhibition of the HEPN ribonuclease RnlA"

### Supplementary Fig. S1

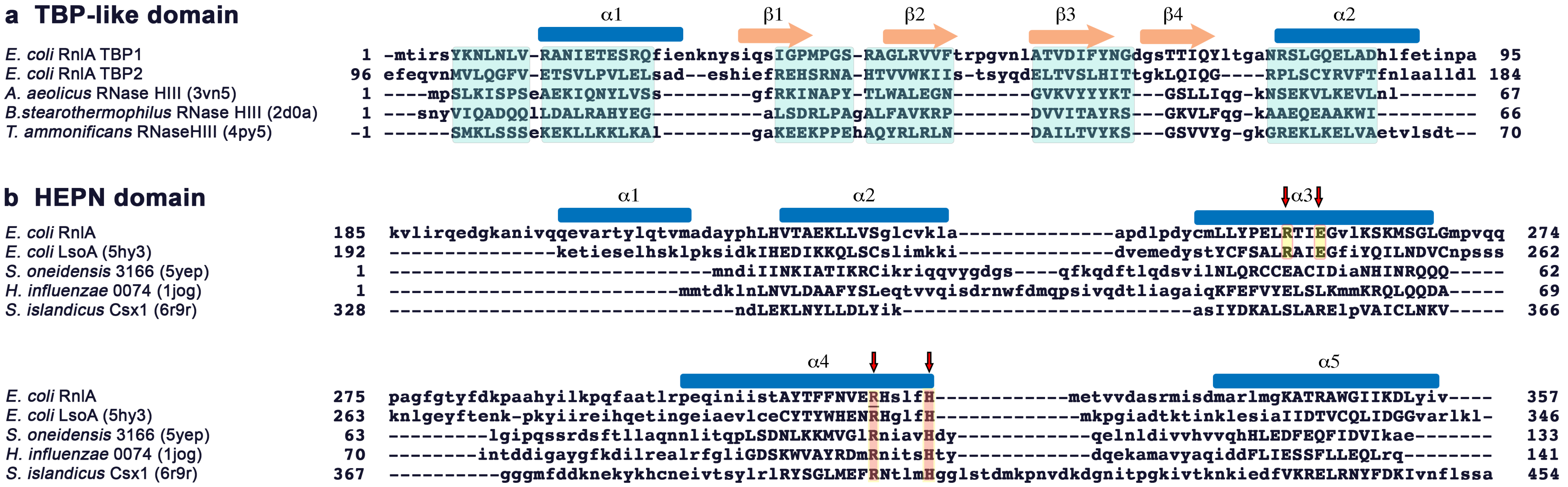


**Supplementary Fig. S1: Structure-based multiple sequence alignment of RnlA domain homologues**. **a** RnlA TBP1 and TBP2 domains are aligned to the TBP-like domains of *Aquifex aeolicus* RNase H3 (PDB ID 3vn5), *Bacillus stearothermophilus* RNase H3 (PDB ID 2d0a) and *Thermovibrio ammonificans* RNase H3 (PDB ID 4py5). **b** Structurally aligned sequences of HEPN domains from HEPN protein representatives from different families: *Escherichia coli* RnlA, *Escherichia coli* LsoA (PDB ID 5hy3), *Shewanella oneidensis* SO_3166 (PDB ID 5yep), *Haemophilus influenzae* HI0074 (PDB ID 1jog), and *Sulfolobus islandicus* Csx1 (PDB ID 6r9r). The alignment was performed by the program POSA[47]. Structurally equivalent residues appear in capital letters and boxed in pale blue for the TBP-like domain. The putative catalytic residues in RnlA are marked by arrows. The arginine and histidine residues in the conserved sequence motif $HX_{4-6}H$ are boxed in red. The arginine and glutamic residues conserved in the RnlA/LsoA HEPN subfamily are boxed in yellow. Secondary structure elements for RnlA are represented on top and labeled; blue bars stand for $\alpha$-helices and pale orange arrows represent $\beta$-strands.

**Supplementary Fig. S2**

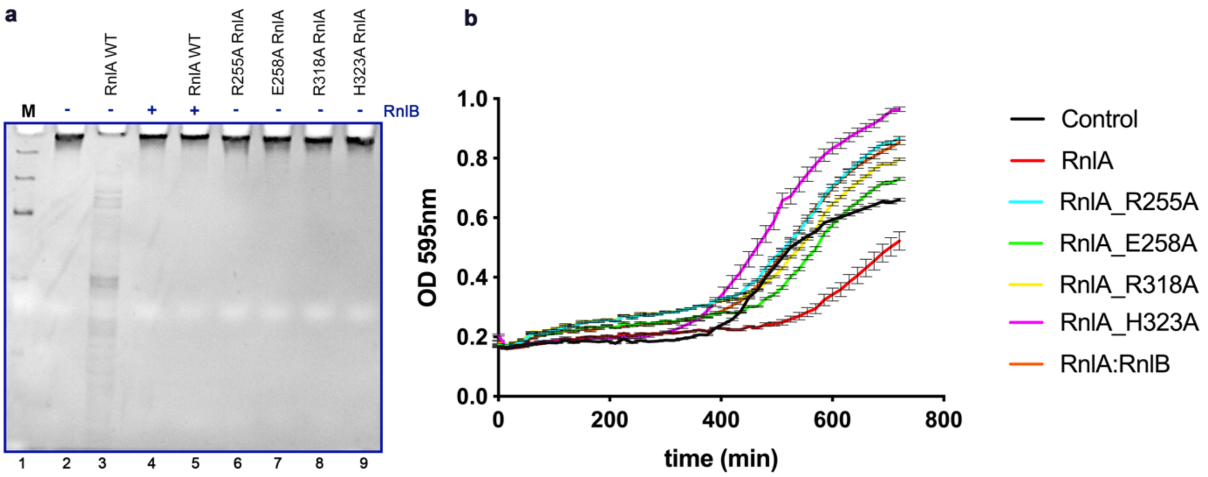


**Supplementary Fig. S2: a RNase Activity and b *in vivo* toxicity of RnlA and single alanine RnlA mutants. a** 5 μM of RnlA and its mutants were incubated with MS2 RNA for 1h prior to denaturing gel electrophoresis. The gel was stained with Ethidium Bromide. 10 μM of RnlB were incubated with RnlA for 10 minutes before the addition of MS2 RNA. Control reactions without protein and with RnlB were also included. M marks the ssRNA standards. Lane 2 corresponds to the untreated RNA substrate. **b** BL21 (DE3) *E. coli* cells containing *pET28a* plasmids with wild-type *rnlA* (n = 24), *R255A_rnlA* (n = 20), *E258A_rnlA* (n = 20), *R318A_rnlA* (n = 20), *H323A_rnlA* (n = 20), and the *rnlAB operon (n = 20)*, were incubated in the presence of IPTG and OD (595 nm) was monitored every 15 minutes. Cells transformed with *pET28* (empty vector, n = 18) were included as a control. Data represent mean ± SEM originating from N > 3 biological repeats.

**Supplementary Fig. S3**

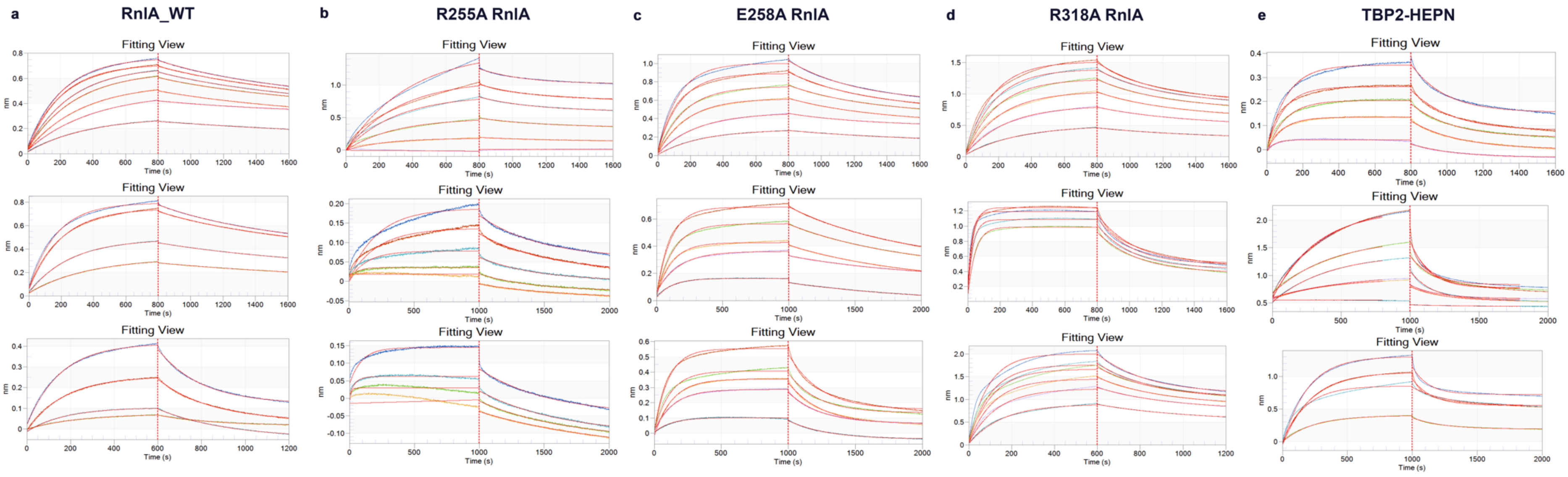


**Supplementary Figure S3. Binding of RnlA, RnlAΔ1-91 (TBP2-HEPN) variant and RnlA single alanine mutants to ssDNA. a** Triplicate binding curves obtained from BLI experiments with ssDNA immobilized on streptavidin sensors. Data were collected in an Octet RED96 instrument and analyzed with the manufacturer data analysis tool v. 9.0 (FORTÉBIO). $K_D$ values are derived from the steady state analysis of the data fits to a 1:1 model, $R_{eq}$ values are extrapolated from the fits to this model. Fitting of the binding curves during association and dissociation to a 1:1 model performed with the Octet Data Analysis software v. 9.0 (FORTÉBIO) for Wild-type RnlA (**a**), R255A RnlA (**b**) E258A RnlA (**c**), R318A RnlA (**d**) and RnlAΔ1-91 **(**TBP2-HEPN) (**e**).

**Supplementary Fig. S4**

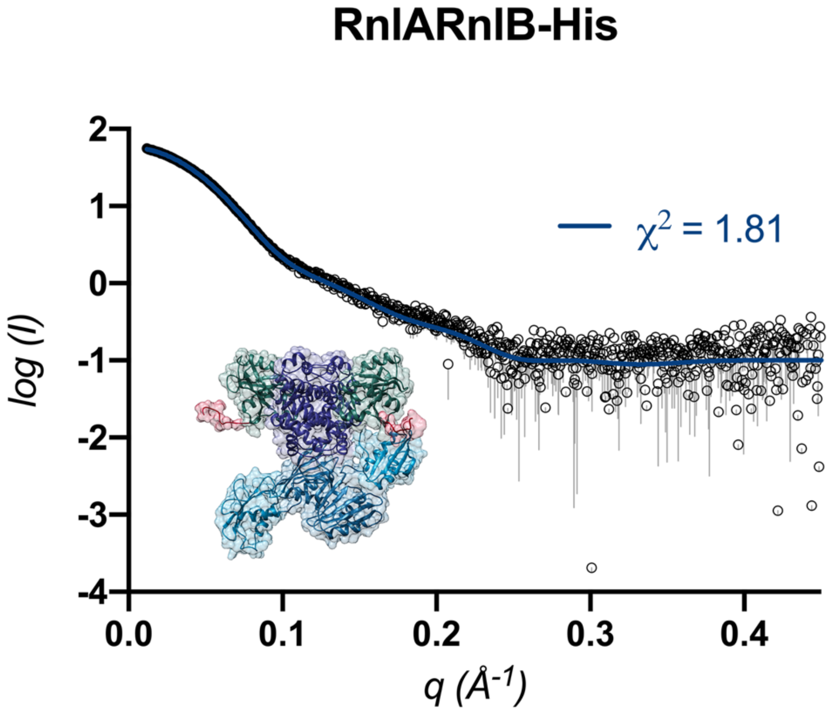


**Supplementary Fig. S4: RnlA:RnlB conformation in solution.** Small angle X-Ray Scattering (SAXS) measurements of the RnlA:RnlB complex. The comparison of the experimental data (open circles, SD are represented as gray lines below the circles) to the theoretical profile for the corresponding all-atom model obtained by the program MODELLER is shown and the corresponding $\chi^2$ value is annotated.

**Supplementary Fig. S5**

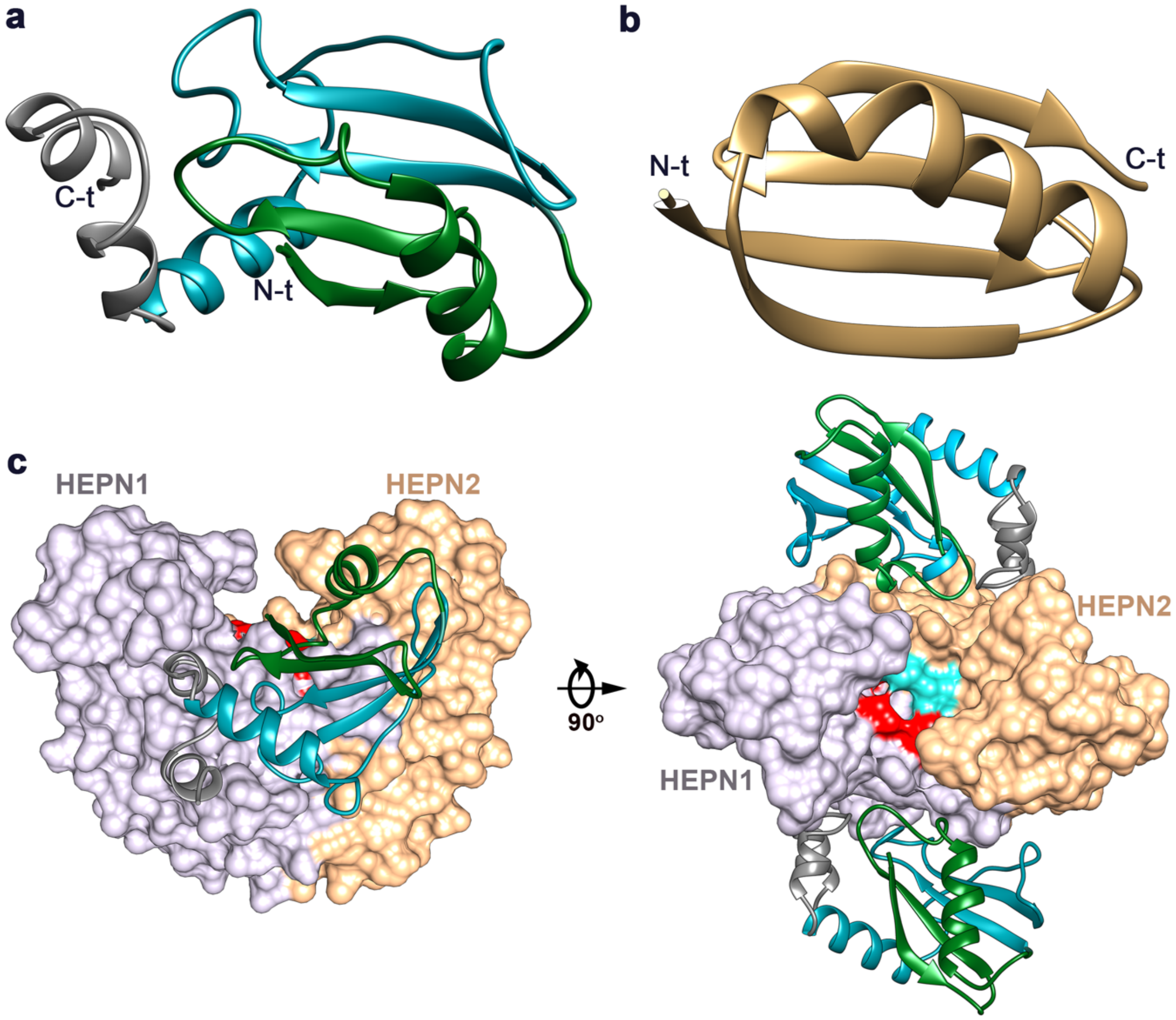


**Supplementary Fig. S5: RnlB docks on both sides of the active site of RnlA, blocking access of the RNA substrate**. **a** Ribbon representation of the RnlB antitoxin. The two ribbon-ribbon-helix motifs are colored in green and cyan respectively and the additional C-terminal short □-helices are colored in grey. The N and C termini are indicated. **b** Ribbon representation of Dmd antitoxin colored in beige with the N and C termini indicated. **c** Interaction between the HEPN domain dimer (surface representation) of RnlA with two RnlB molecules (ribbon representation). The active site residues of RnlA (R255, E258, R318, H323) are highlighted in cyan and red, respectively, for the two HEPN subunits. RnlB does not contact the active site residues of RnlA, which are partially buried in the canonical HEPN dimer interface. Two views rotated by 90° along the *x* axis are shown.

**Supplementary Fig. S6**

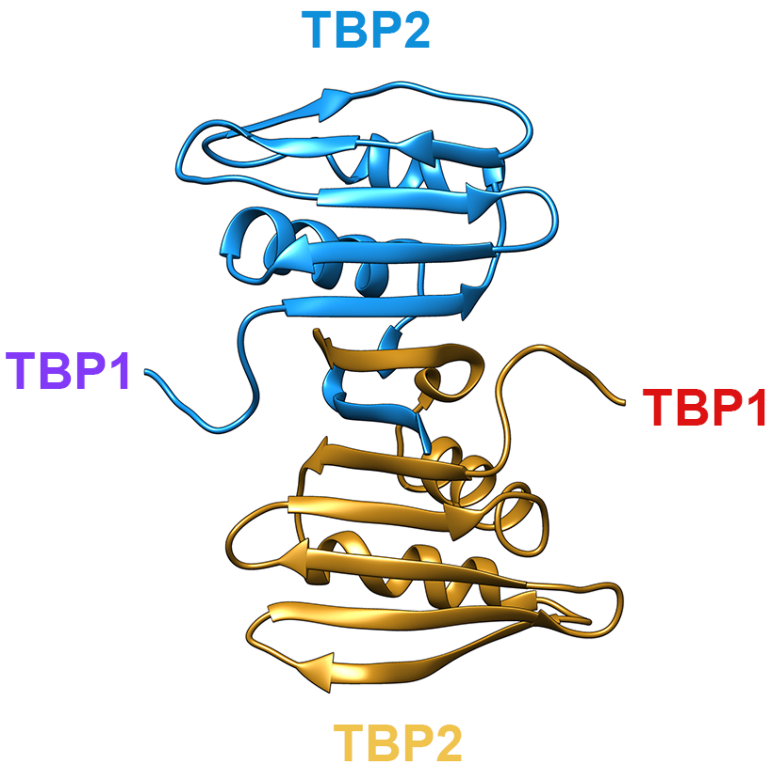


**Supplementary Fig. S6: RnlA TBP2 dimerization in RnlB-induced RnlA dimer.** Ribbon representation of TBP2 dimerization shown from the top. TBP1 and HEPN domains are not shown for clarity. The positions of TBP1 toward the N-terminus are indicated. RnlA domains are colored differently in each chain, A: TBP2 in yellow, B: TBP2 in light blue.

Supplementary Fig. S7

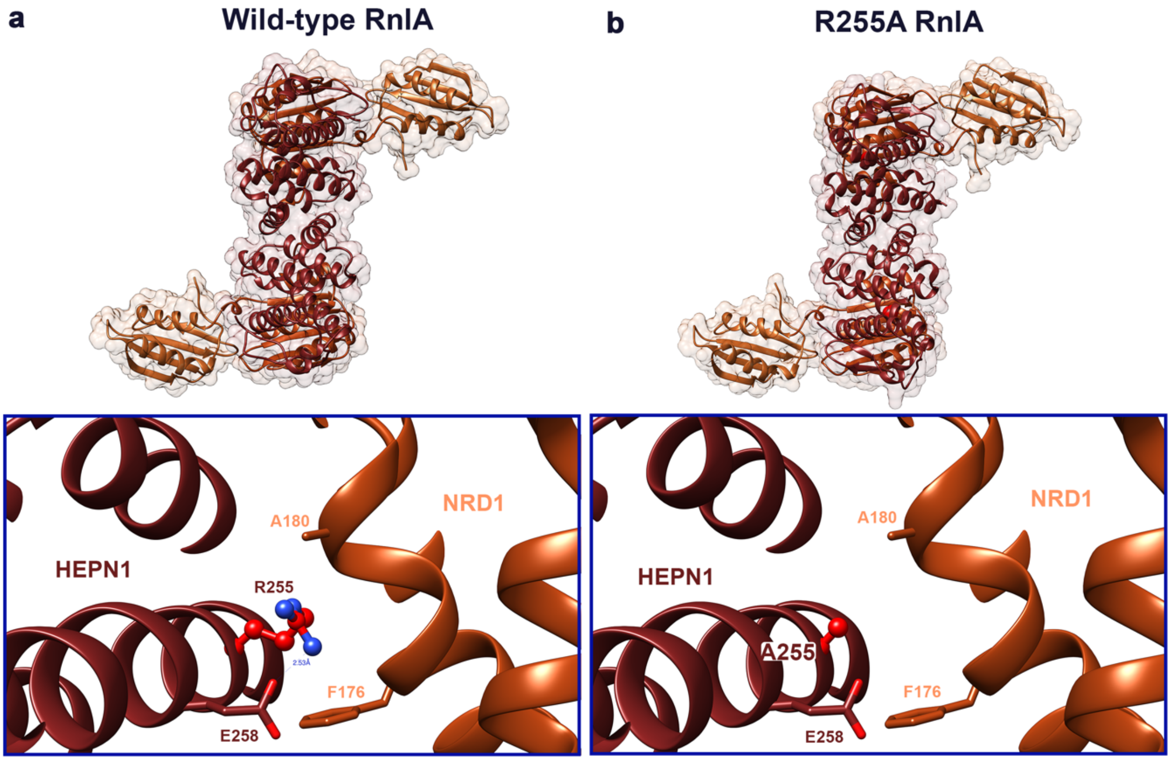


**Supplementary Fig. S7: R255A RnlA mutant exists as the free dimer in the crystal**. The crystallographic structures of wild-type RnlA (**a**) and R255A RnlA (**b**). Stick and ball representation of R255 in free RnlA (**a**) and R255A mutation (**b**). Residues contacting R255 are represented as sticks and chains are represented as ribbons.

**Supplementary Fig. S8**

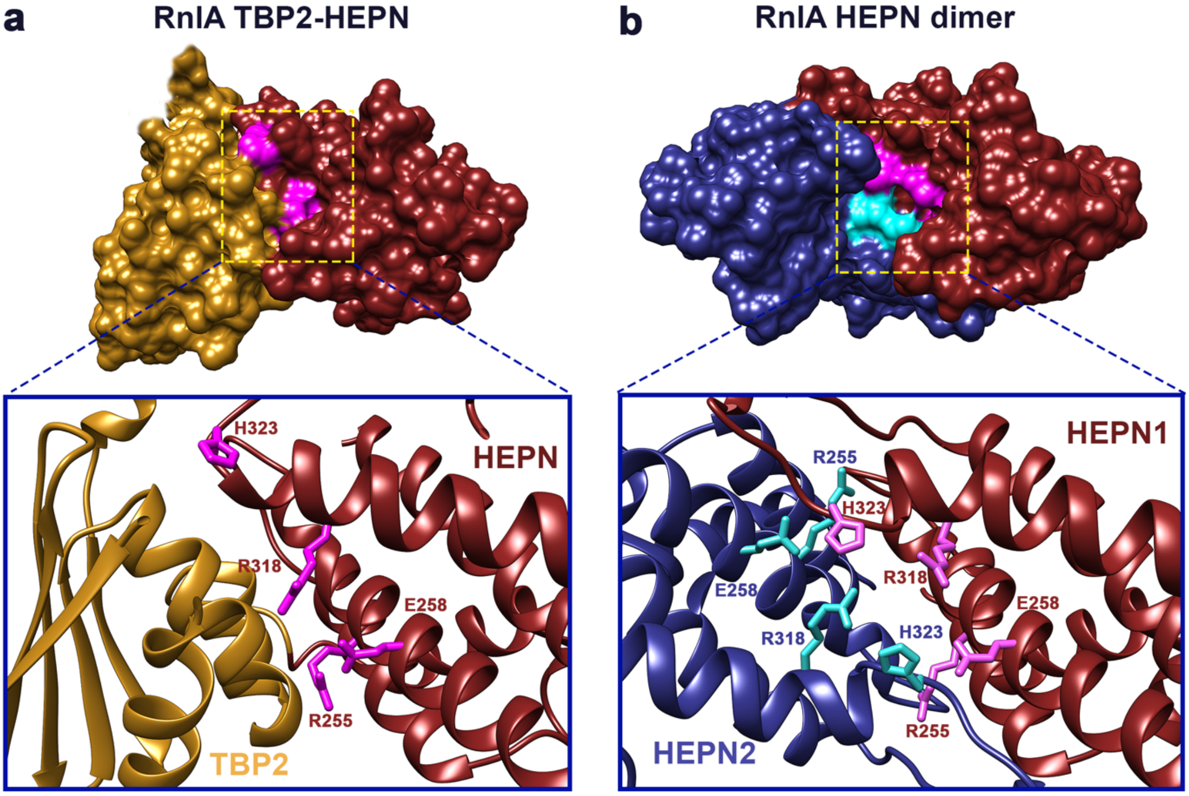


**Supplementary Fig. S8: RnlA catalytic residues in the TBP2-HEPN and HEPN-HEPN interfaces. a** Surface representation of the HEPN (dark red) and TBP2 (yellow) domains in chain A of RnlA. The TBP1 domain has been removed for clarity. The surface of the catalytic residues is colored magenta. In the zoom-in, the positions of R255, E258, R318 and H323 are shown in stick representation. They are buried on the interface with the TBP2 domain and both domains have to move relative to each other to make the putative active site accessible to the RNA substrate. **b** Surface representation of the canonical RnlA HEPN-dimer as seen in the RnlA:RnlB complex. One HEPN domain is colored dark red and is in the same orientation as the HEPN domain in **a**. Its active site is colored magenta. The second HEPN domain is colored dark blue and its active site cyan. The zoom-in shows a stick representation of the catalytic residues R255, E258, R318 and H323 in both domains (ribbon representation). Both sets of active sire residues interact with each other and are partially buried.

**Supplementary Fig. S9**

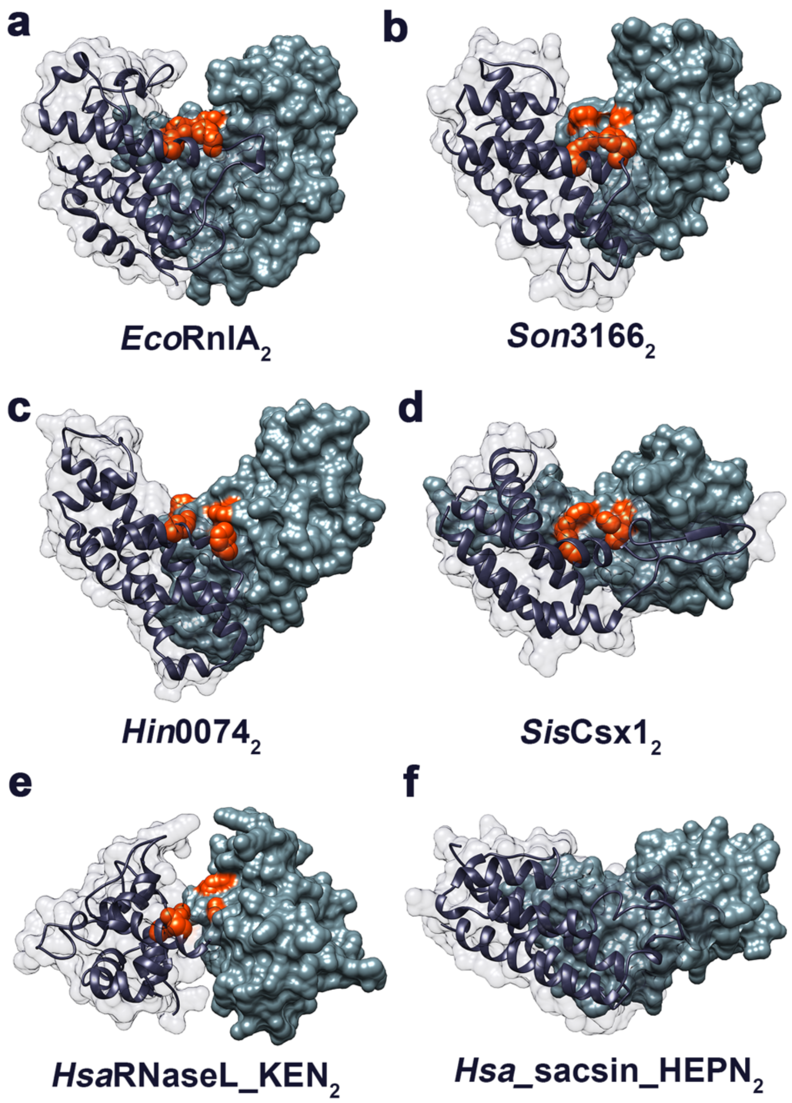


**Supplementary Fig. S9: Comparison of HEPN-domain dimerization across this superfamily.** One representative is shown for each of the six different HEPN families. **a** The canonical HEPN-domain dimer of RnlA as observed in the RnlA:RnlB complex. One chain is shown in dark blue ribbon with a transparent surface, the other chain as a solid dark green surface. **b** HEPN dimer of SO_3166 from *Shewanella oneidensis* (PDB ID 5yep) as seen in the SO_3166-SO_3165 TA complex. Here the putative active site residues are directly covered by the antitoxin. **c** Hi0074 from *Haemophilus influenza* (PDB ID 1jog) which is a putative nucleotidyltransferase. **d** SisCsx1 CRISPR-Cas RNase from *Sulfolobus islandicus* (PDB ID 6r9r). **e** KEN domain from human RNase L (PDB ID 4oav). **f** Nucleotide binding HEPN domain dimer from human Sacsin (PDB ID 3o10). This domain has no RNase activity and lacks the characteristic $RX_{4-6}H$ motif, which is represented as red spheres in all other HEPN proteins.

**Supplementary Fig. S10**

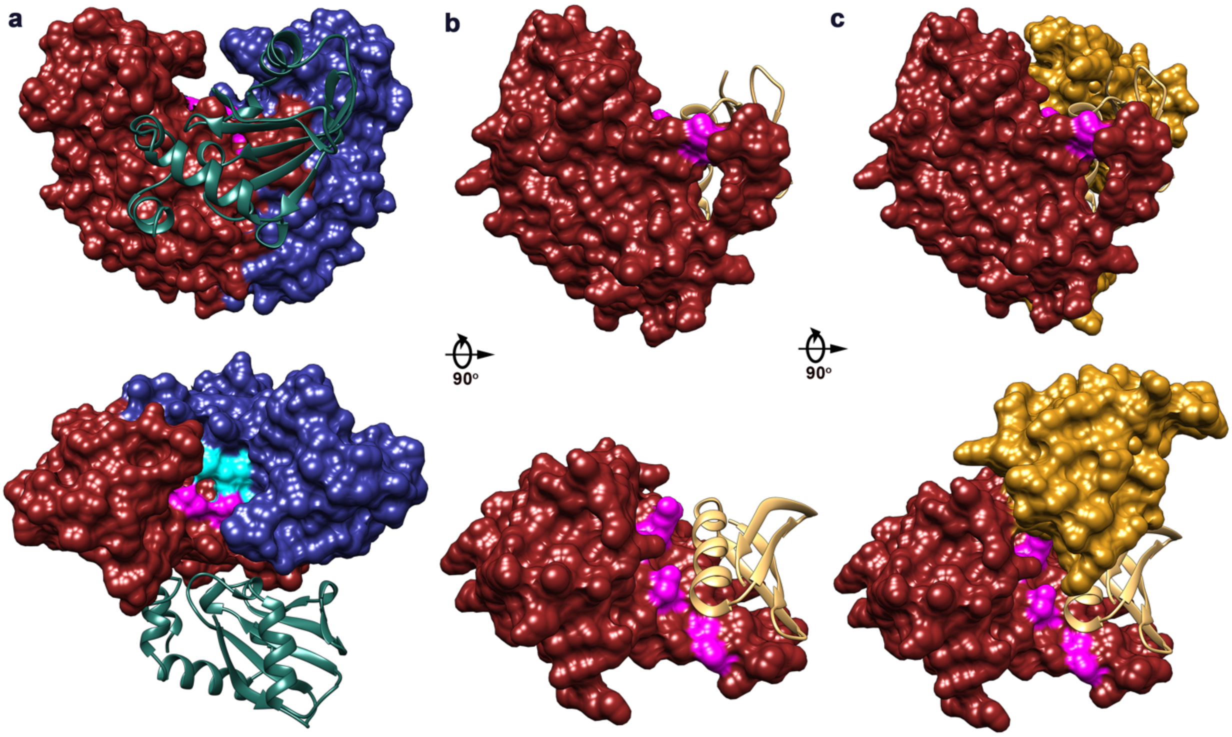


**Supplementary Fig. S10: Comparison of the inhibitory mechanism of RnlB and Dmd antitoxins.** Ribbon representation of **a** RnlB monomer (dark green) and **b** Dmd antitoxin monomer (beige) bound to RnlA HEPN-dimer (HEPN1 in surface representation colored dark red and HEPN2 in surface representation colored dark blue), and LsoA HEPN domain (surface representation in dark red), respectively. RnlA HEPN1 and LsoA HEPN are superposed and two 90°-related orientations along the *x* axis are shown. **c** Ribbon representation of Dmd (beige) bound to LsoA with TBP2 shown as yellow surface.

### Supplementary Tables

#### Supplementary Table S1. Crystallographic data collection and refinement statistics

| | RnlA:RnlB | RnlA | RnlAΔ1-91 | R255A RnlA |
|---|---|---|---|---|
| **Wavelength (Å)** | 0.9801 | 0.9724 | 0.9786 | 0.9801 |
| **Resolution range (Å)** | 48.78 - 2.64 (2.73 - 2.64) | 47.9 - 2.99 (3.09 - 2.99) | 45.19 - 1.94 (2.01 - 1.94) | 45.85 - 3.89 (4.03 - 3.89) |
| **Space group** | C 1 2 1 | P 21 21 21 | P 21 21 2 | P 21 21 21 |
| **Unit cell** | 243.32 133.58 55.64 90° 95.11° 90° | 64.32 100.8 154.09 90° 90° 90° | 69.65 81.67 54.25 90° 90° 90° | 64.23 104.93 152.94 90° 90° 90° |
| **Total reflections** | 350150 (31175) | 144872 (12457) | 305397 (24063) | 100831 (9527) |
| **Unique reflections** | 51657 (4871) | 20825 (1869) | 23246 (1700) | 9959 (977) |
| **Multiplicity** | 6.8 (6.4) | 7.0 (6.4) | 13.1 (11.1) | 10.1 (9.8) |
| **Completeness (%)** | 99.24 (93.63) | 98.32 (90.42) | 96.47 (74.17) | 99.61 (98.29) |
| **Mean I/sigma(I)** | 12.10 (1.51) | 9.36 (1.12) | 12.72 (0.84) | 4.44 (0.88) |
| **Wilson B-factor** | 70.41 | 81.74 | 33.50 | 124.23 |
| **R-merge** | 0.11 (0.81) | 0.17 (1.74) | 0.12 (2.36) | 0.41 (2.31) |
| **R-meas** | 0.119 (0.88) | 0.18 (1.9) | 0.13 (2.47) | 0.43 (2.43) |
| **R-pim** | 0.045 (0.34) | 0.07 (0.73) | 0.035 (0.71) | 0.13 (0.76) |
| **CC1/2** | 0.99 (0.78) | 0.99 (0.50) | 0.99 (0.46) | 0.98 (0.54) |
| **CC*** | 0.99 (0.93) | 0.99 (0.82) | 1 (0.79) | 0.99 (0.84) |
| **Reflections used in refinement** | 51586 (4821) | 20809 (1869) | 22523 (1683) | 9946 (976) |
| **Reflections used for R-free** | 2576 (239) | 1041 (94) | 1997 (148) | 497 (48) |
| **R-work** | 0.19 (0.28) | 0.20 (0.34) | 0.19 (0.31) | 0.21 (0.33) |
| **R-free** | 0.24 (0.28) | 0.25 (0.36) | 0.23 (0.35) | 0.30 (0.33) |
| **CC (work)** | 0.95 (0.83) | 0.94 (0.70) | 0.95 (0.61) | 0.96 (0.71) |
| **CC (free)** | 0.94 (0.80) | 0.89 (0.70) | 0.93 (0.53) | 0.923 (0.67) |
| **Number of non-hydrogen atoms** | 7065 | 5380 | 4544 | 5394 |
| **macromolecules** | 7015 | 5175 | 4216 | 5394 |
| **ligands** | | 2 | 328 | - |
| **solvent** | 50 | 3 | 534 | - |
| **Protein residues** | 943 | 692 | 0.008 | 689 |
| **RMS (bonds)** | 0.012 | 0.007 | 1.22 | 0.005 |
| **RMS (angles)** | 1.48 | 1.31 | 97.36 | 1.08 |
| **Ramachandran favored (%)** | 94.53 | 95.76 | 2.64 | 92.07 |
| **Ramachandran allowed (%)** | 5.26 | 4.24 | 0.00 | 7.20 |
| **Ramachandran outliers (%)** | 0.21 | 0.00 | 0.00 | 0.73 |
| **Rotamer outliers (%)** | 0.72 | 1.05 | 3.66 | 1.03 |
| **Clashscore** | 10.09 | 8.70 | 38.77 | 5.66 |
| **Average B-factor** | 96.02 | 81.74 | 38.45 | 137.15 |
| **macromolecules** | 96.22 | 81.76 | 42.98 | 137.15 |
| **ligands** | | 75.38 | 4216 | - |
| **solvent** | 67.18 | 54.87 | 328 | - |
| **Number of TLS groups** | 28 | 2 | 1 | 6 |

*Statistics for the highest-resolution shell are shown in parentheses.*

### Supplementary Table S2. Sample details for SAXS measurements

| | RnlA:RnlB | RnlA | R255A RnlA | RnlAΔ1-91 | D245R RnlA |
|---|---|---|---|---|---|
| **SASBDB.org accession code** | SASDHW7 | SASDHX7 | SASDHY7 | SASDKL2 | SASDKS9 |
| **Organism** | *Escherichia coli K-12* | | | | |
| **Source** | *Escherichia coli* | | | | |
| **Uniprot ID** | P52129, P52130 | P52129 | P52129 | P52129 | P52129 |
| **Ext. Coeff (A280, 0.1 % (w/v))** | 0.734 | 0.755 | 0.797 | 0.833 | 0.794 |
| **MW (Da)** | 109 918 | 84430 | 83 685 | 62 200 | 83 938 |
| **SEC-SAXS column** | Shodex KW404-4F 500 kDa | batch | Shodex KW404-4F 500 kDa | Agilent Bio-SEC 3 | batch |
| **Loading concentration (mg ml$^{-1}$)** | 5.9 | 2.4 | 13 | 24 | 0.34 – 2.2 |
| **Injection volume (ml)** | 40 | 40 | 40 | 60 | 40 |
| **Flow rate (ml min$^{-1}$)** | 0.2 | - | 0.2 | 0.2 | - |
| **Buffer** | 20 mM Tris pH 8.0, 150 mM NaCl, 1 mM TCEP | | | | |

### Supplementary Table S3. SAXS data collection parameters

| | RnlA:RnlB | RnlA | R255A RnlA | RnlAΔ1-91 | D245R RnlA |
|---|---|---|---|---|---|
| **Beamline** | BM29 ESRF Grenoble | | | Swing SOLEIL | |
| **Detector** | Pilatus 1M | | | EIGER – 4M | |
| **Wavelength (Å)** | 0.9919 | 0.9919 | 0.9919 | 1.0331 | 1.0331 |
| ***q* measurement range (Å$^{-1}$)** | 0.011 - 0.481 | 0.015 – 0.404 | 0.010 – 0.428 | 0.003 – 0.45 | 0.008 – 0.400 |
| **Sample temperature (°C)** | 20 | | | 16 | 16 |
| **Sample configuration** | SEC-SAXS | Batch mode | SEC-SAXS | SEC-SAXS | Batch mode |
| **Frames averaged** | 90-130 | | 110-160 | 231-281 | |

### Supplementary Table S4. SAXS structural parameters

| | RnlA:RnlB | RnlA | R255A RnlA | RnlAΔ1-91 | D245R RnlA |
|---|---|---|---|---|---|
| **Guinier analysis** | | | | | |
| **I(0) (AU)** | 57.94 ± 0.07 | 49.96 ± 0.12 | 86.95 ± 0.12 | 0.078 ± 0.00002 | 0.052 ± 0.002 |
| **$R_g$ (Å)** | 34.70 ± 0.29 | 37.29 ± 0.19 | 36.83 ± 0.28 | 30.05 ± 0.06 | 42.61 ± 1.27 |
| **$q_{min}$ (Å$^{-1}$)** | 0.016 | 0.015 | 0.014 | 0.011 | 0.008 |
| **Coefficient correlation, $R^2$** | 0.94 | 0.97 | 0.95 | 0.90 | 0.98 |
| **M (Da) from I(0)** | 94 225 | 85 650 | 78 500 | 52 754 | 91 175 |
| **Expected M (Da)** | 109 972 | 83 874 | 83 685 | 31 109 | 83 740 |
| ***P(r)* analysis** | | | | | |
| **I(0) (AU)** | 56.63 | 48.68 | 89.24 | 0.08 | 0.05 |
| **$R_g$ (Å)** | 33.40 | 36.02 | 40.89 | 30.22 | 45.58 |
| **$D_{max}$ (Å)** | 100.4 | 107.3 | 201.2 | 104.3 | 159.1 |
| ***q* range (Å$^{-1}$)** | 0.003 - 0.395 | 0.001-0.295 | 0.002 – 0.218 | 0.011 – 0.291 | 0.01 – 0.18 |
| **Porod Volume (Å$^3$)** | 161 657 | 134 012 | 133 054 | 89 682 | 177 741 |

### Supplementary Table S5. SAXS Atomistic modeling

| | RnlA:RnlB | RnlA | R255A RnlA | RnlAΔ1-91 | D245R RnlA |
|---|---|---|---|---|---|
| **Crystal Structures (PDB IDs)** | 6Y2P | 6Y2Q | 6Y2R | 7AEX | Based on 6Y2Q |

| **MODELLER** | RnlB C-terminal His-tag | (N-terminal His-tag) Free dimer | (N-terminal His-tag) RnlB-induced RnlA-dimer | (N-terminal His-tag) Free RnlA-like | (N-terminal His-tag) RnlB-induced RnlA-like | (N-terminal His-tag) Free RnlA-like (TBP1 truncated) | (N-terminal His-tag) RnlB-induced RnlA-like (TBP1 truncated) | (N-terminal His-tag) Free dimer | (N-terminal His-tag) RnlB-induced RnlA-dimer |
|---|---|---|---|---|---|---|---|---|---|
| **MultiFoXS** | | | | | | | | | |
| $\chi^2$ | 1.81 | 1.06 | 5.06 | 3.39 | 39.99 | 3.10 | 66.16 | 4.1 | 21.5 |
| **Predicted $R_g$ (Å)** | 33.60 | 35.29 | 32.13 | 35.29 | 31.49 | 30.37 | 29.73 | 36.02 | 32.1 |
| $c_1, c_2$ | 1.02, -0.50 | 1.03, -0.50 | 1.05, 1.46 | 1.03, -0.50 | 1.05, 1.08 | 1.03, -0.45 | 1.03, 0.40 | 1.04, 0.54 | 1.05, 2.00 |
